# Revising the ortholog conjecture in cross-species comparison of scRNA-seq data

**DOI:** 10.1101/2024.06.21.600109

**Authors:** Yuyao Song, Detlev Arendt, Irene Papatheodorou, Alvis Brazma

## Abstract

The cross-species comparison of expression profiles uncovers functional similarities and differences between cell types and helps refining their evolutionary relationships. Current analysis strategies typically follow the ortholog conjecture, which posits that the expression of orthologous genes is most similar between species. However, the extent to which this holds true at different evolutionary distances is unknown. Here, we systematically explore the ortholog conjecture in comparative scRNA-seq data. We devise a robust analytical framework, GeneSpectra, to classify genes by expression specificity and distribution across cell types. Our analysis reveals that genes expressed ubiquitously across nearly all cell types exhibit strong conservation of this pattern across species, as do genes with high expression specificity. In contrast, genes within intermediate specificity fluctuate between classes. As expected, ortholog expression becomes more divergent with increased species distance. We also find an overall correlation between similarity in expression profiles and sequence conservation. Finally, our results allow identifying gene classes with highest probability of expression pattern conservation that are most useful for cell type alignment between species. Calibrating reliance on the ortholog conjecture for individual genes, we thus provide a comprehensive framework for the comparative analysis of single cell data.

## Introduction

Over millions of years, evolution has shaped the building blocks of life, giving rise to the plethora of species we see today. Comparing gene expression data across species at single-cell resolution with single-cell or single-nucleus RNA-seq (scRNA-seq/snRNA-seq) can shed light on the evolutionary links between cell types ^1–3^. Current computational strategies for cross-species comparisons of scRNA-seq data typically rely on two approaches: expression pattern correlation and data integration ^4–6^. Both approaches require matching genes by orthology to measure their expression similarity between cells across species. To be specific, in correlation analysis, one-to-one (1-2-1) orthologs are often used as the only shared features to calculate the degree of cell type similarity ^7–11^. In data integration, algorithms require orthology-mapped data as input to align transcriptomically similar cell populations ^7,12–14^. For both strategies, the key assumption is that cell types from different species are alike if they have similar relative expression patterns of orthologs.

This assumption underpinning computational cross-species scRNA-seq comparisons is rooted in the hypothesis known as the *ortholog conjecture* ^15–17^. In an evolutionary sense, the hypothesis posits that orthologs typically perform analogous functions in the respective organisms ^17,18^. The conjecture further proposes for function predictions that orthologs are superior predictors due to greater functional conservation than within-species paralogs (in-paralogs) ^19^. As cell type gene expression patterns can be considered as a functional readout of the gene, the hypothesis had been implicitly transferred to the analysis of cross-species scRNA-seq data. According to the ortholog conjecture, orthologs are expected to exhibit similar cell type expression patterns across species, implying that their expression should correlate across corresponding cell types in different organisms and thus can then be used to evaluate the similarity of cell types between species. So far, the conjecture has seen broad application to translate disease models and functional studies from experimental organisms to humans.

It is crucial to point out that the validity of the ortholog conjecture has been intensely debated. Literature focus on evaluating the degree to which orthologs were similar under certain functional measurements, and whether the similarity between orthologs was higher than that of paralogs ^15,16,19–24^. It is generally agreed that having an empirical, quantitative estimate of the functional similarity of orthologs, respective to their evolutionary relationship, is a more rigorous approach ^15,25^. Therefore it is important to develop methods for measuring and establish a baseline for the extent to which the ortholog conjecture applies in terms of cell type expression patterns in scRNA-seq data.

Several inherent features of scRNA-seq data and common analytical choices pose new challenges to assessing the ortholog conjecture. In addition to technical while mitigatable issues such as data sparsity and heteroscedasticity ^26^, there is a more fundamental problem, which we denote as the *challenge of circularity* (Figure 1a). This challenge arise when gene expression data are first used to define highly-granular cell types and mapping them between species, while subsequently, the same expression data were used to assess the ortholog conjecture. In other words, the difficulty is that on a highly fine-grained level, cell types are often only defined transcriptomically and then these cell type assignments are transferred cross species based on the similarity in expression patterns of 1-2-1 orthologs. But if so, then assessing the ortholog conjecture based on the same or similar data inevitably introduces circularity, which means that the perceived degree of gene expression conservation is artificially inflated. This challenge is currently difficult to overcome, but it can be mitigated when the cell type annotation is biologically robust and supported by biological evidence from other modalities beyond gene expression. As long as the annotations maintain confidence at the cell type or cell class level, the results will reflect genuine biological patterns at least to a large degree. Therefore, it is necessary to carefully determine what kind of cell type annotation is suitable for the particular cross-species single-cell gene expression pattern analysis. Ideally, the datasets from different species should be annotated independently with appropriate cell type granularity.

**Figure 1.**
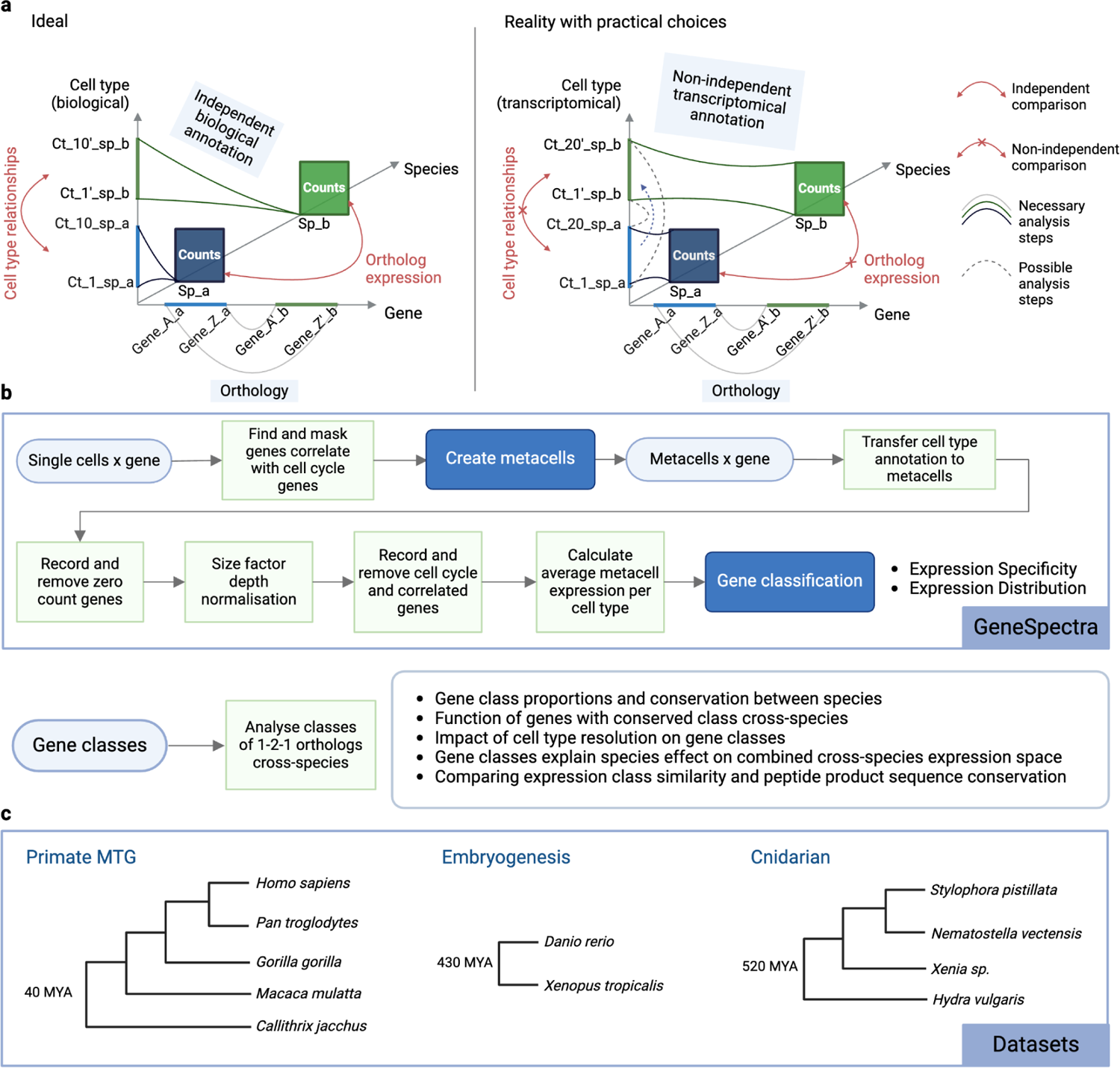
Schematic of the GeneSpectra framework and dataset used in this study. ^64^. (a) A graphical representation of an ideal analysis scenario vs.current practical strategies for cross-species scRNA-seq data analysis. To achieve truly independent comparisons of cell types and ortholog expression between species, cell types are annotated independently between species, rather than solely based on gene expression. In reality, cell types are often transcriptomically defined, and potentially transferred between species via orthologous genes. This could inflate the perceived similarity between cell types as well as the ortholog expression patterns. (b) Workflow of the GeneSpectra computational framework. (c) Species tree of datasets used in this study. Counts, the scRNA-seq gene expression count table; 1-2-1, one-to-one; OG, ortholog groups; MTG, middle temporal gyrus; MYA, million years ago.

For comparing evolutionary distant species, the reliance on the ortholog conjecture hypothesis becomes even less reliable. First, there are few 1-2-1 orthologs between distant species genomes, and the gene orthology relationship can be obscure ^27^. Secondly, the genome context has diverged, and tissues experience concerted expression evolution which leads to correlated expression changes between cell types in that tissue ^1,28^. Lastly, new genes that emerged in a species can play a major role in the species’ cell type expression landscape and contribute to species-unique cell populations ^29–31^. Several recent cross-species integration tools have been developed to tackle the restriction by gene orthology in order to improve performance among remote species ^32–35^. However, we here argue that the fundamental challenge lies not in the development of additional algorithms but in our limited understanding of which measurable expression features are conserved over such long evolutionary timescales.

Here we show that the degree of the applicability of the ortholog conjecture depends not only on the evolutionary distance between the species, but also on the specific expression patterns of the respective genes. In like manners to the Human Protein Atlas (HPA) project ^36,37^, we classify genes by their cell type expression specificity and distribution, then we compare the classes between orthologs. We show that the genes that are expressed ubiquitously or non-specifically across nearly all cell types exhibit strong conservation of this expression pattern across species at different evolutionary distances. Similarly, genes with high expression specificity in one cell type or cell type family also retain their profiles, albeit to a lesser extent. The expression patterns of the genes with intermediate specificity are the least conserved. As expected, with the increase of species evolutionary distance, ortholog expression patterns become generally more divergent. We also show an overall positive relationship between expression pattern similarity and protein sequence conservation across different types of orthologs. To facilitate reproducibility and future work, we make available an open source tool called GeneSpectra (https://github.com/Papatheodorou-Group/GeneSpectra), as well as the expression patterns of all genes used in our analysis.

## Results

### Robust framework for gene classification by cell type expression specificity and distribution

To describe the cell type pattern of gene expression and examine their conservation between orthologs, we first performed a gene classification analysis (Figure 1b). The classification is based on the cell type specificity and distribution of the expression of genes, with criteria derived from the HPA classes ^36^ (Table 1). We developed an open-source Python module GeneSpectra to streamline the analysis (freely available at https://github.com/Papatheodorou-Group/GeneSpectra). The GeneSpectra module first reduces the sparsity of scRNA-seq data, then, after normalisation, applies gene classification. It further provides functionalities to compare gene class conservation between orthologs, utilizing results from the ortholog mapping method of choice.

**Table 1.** Gene expression specificity and distribution classification. The gene classes and criteria are modified from the Human Protein Atlas ^36^ gene classification on single-cell RNA-seq data. CPM, counts per million.

| Gene expression specificity | Class | Criteria |
| --- | --- | --- |
|  | Lowly expressed | Average expression in all cell type < 1 CPM |
|  | Low cell type specificity | All genes which do not satisfy the criteria of being specific |
| | Group Enhanced | Expression level (CPM) in some cell types ≥ 4-fold average expression across all cell types |
| | Cell type Enhanced | Expression level (CPM) in one cell type ≥ 4-fold average expression across all cell types |
| | Group enriched | Average expression (CPM) in a group of cell types ≥ 4-fold maximum expression in all other cell types |
| | Cell type enriched | Expression (CPM) in one cell type ≥ 4-fold maximum expression in all other cell types |
| Gene expression distribution | Class | Criteria |
|  | Lowly expressed | Average expression in all cell type < 1 CPM |
|  | Expressed in single | Expression level over 1 CPM in one cell type |
|  | Expressed in few | Expression level over 1 CPM in less than 30% of all cell types |
|  | Expressed in many | Expression level over 1 CPM in over 30% but less than 90% of all cell types |
|  | Expressed in most | Expression level over 1 CPM in over 90% of all cell types |

Raw scRNA-seq data represent repeated sparse sampling of biological single cells due to the limited transcript capturing in scRNA-seq experiments ^38^. To obtain a sparsity-corrected cell type expression profile for each gene, without disrupting the single-cell granularity, we employed the metacell approach ^39,40^ (Supplementary figure 1, see details in Methods). The metacell approach groups together single cells to provide a more robust estimation of transcription states, which is essential for computing statistics such as mean expression ^40^. We calculated the averaged expression data among metacells of the same type as the cell type expression profile input to GeneSpectra.

Following size factor normalisation, we performed the gene classification using annotated cell types for each species. The gene classification criteria were derived from the HPA classes which were used to describe the tissue patterns of gene expression (Table 1). Genes were grouped by their expression specificity, a relative criteria describing how elevated is the gene expression in certain cell types. In practice, enriched genes exhibit expression levels at least 4-fold by default higher in certain cell types or groups compared to the second-highest expression level, while enhanced genes show expression levels at least 4-fold higher than the average level across all cell types. Meanwhile, genes can also be classified by their expression distribution which is an absolute criterion for how widely the gene is expressed across cell types. Lastly, genes showing no cell type specificity or have low expression level have their respective classes.

We mainly applied the GeneSpectra classification to 3 datasets: a middle temporal gyrus (MTG) dataset from 5 primates ^7^; an embryogenesis dataset from zebrafish and xenopus ^41,42^ and a dataset consisting of whole-body sequencing of 4 Cnidarian species ^43^ (hereafter refer to as primate MTG; embryogenesis and Cnidarian datasets, respectively, Figure 1c).

We applied GeneSpectra to the primate MTG dataset using the “subclass” cell type labels from the original publication. The analysis showed that the specificity and distribution class proportions for all genes that passed minimum expression for quantification were highly similar among the species, with old World monkeys (marmoset, macaque) and great apes (chimpanzees, gorilla) showing a notable resemblance within their respective groups (Figure 2a, b). Notably, around 40%-50% of genes showed low cell type specificity or expression over 90% of cell types, which was consistent with the percentage of “housekeeping” genes (44%) found across tissues in the HPA ^37^. Comparing the percentages of genes showing enrichment/enhancement between cell types suggested that each of the non-neuronal cell types had a relatively higher number of specific genes than per neuron cell type, and the percentages stayed consistent across species (Figure 2c, d). We further observed that transcriptomically similar cell types shared more group specific genes, with non-neuronal cells again showing a high number of intersections (Supplementary Figure 2).

**Figure 2.**
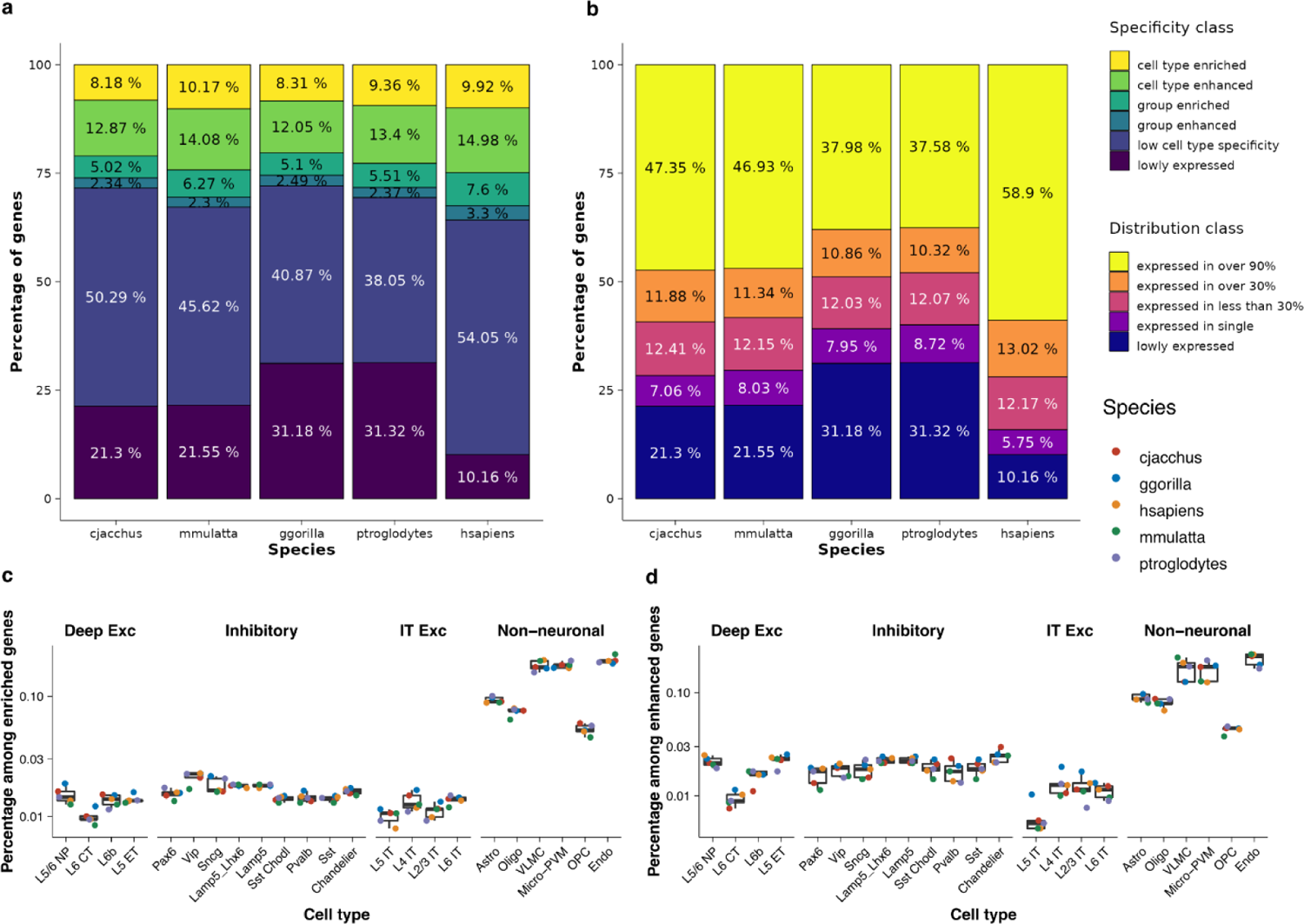
Gene class distribution in the primate MTG datasets. (a) Gene specificity class and (b) gene distribution class percentage of all genes in each species. (c) percentage of enriched genes and (d) enhanced genes in each cell type. Data points were shown as dots. Note that y axis is at log10 scale in (c) and (d). The bar in the boxplot shows the arithmetic mean, lower and upper hinges correspond to the first and third quartiles, whiskers extend from the hinge to the largest value no further than 1.5 * interquartile range.

### Gene class divergence vary among 1-2-1 orthologs with different cell type expression patterns

Since 1-2-1 orthologs were typically considered as matching features in cross-species expression analysis, we asked to what degree they align in terms of cell type expression specificity and distribution. Therefore, we examined if 1-2-1 orthologous genes stayed in the same class between pairs of species in all 3 datasets. In this section, we focus on the primate MTG dataset to describe patterns of the gene class conservation in closely related species.

We observed a notable frequency of specificity class switching among enriched or enhanced genes across species, with lowly specific genes predominantly maintaining their low specificity. Cell type enriched genes appeared to be slightly more conserved than group enriched or enhanced genes. Figure 3a shows the average gene specificity class conservation matrix of 1-2-1 orthologs in pairs of species for the primate MTG dataset (see Methods for matrix calculation, see Supplementary figure 3 for each species pair results).

**Figure 3.**
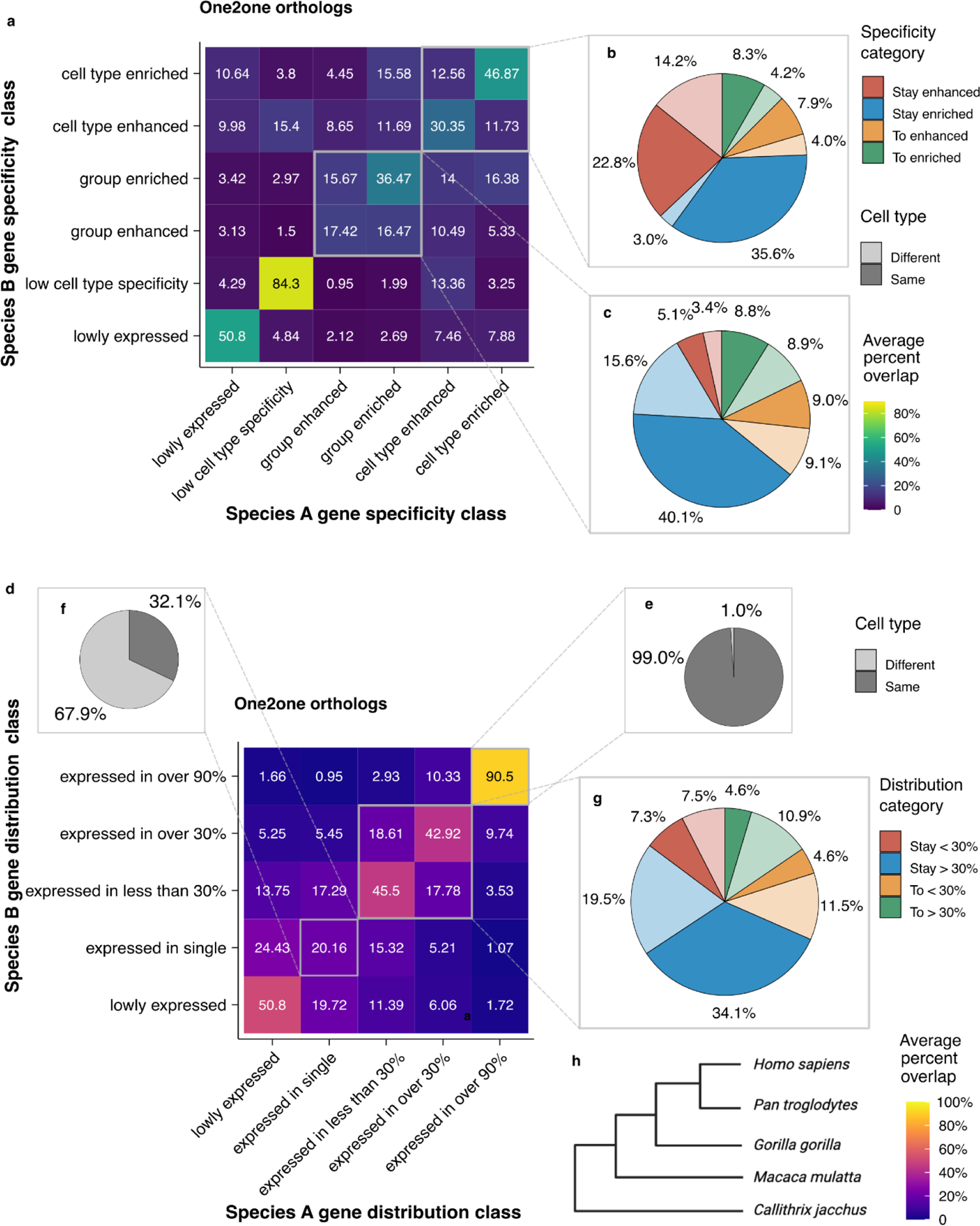
Gene class conservation among 1-2-1 orthologs in primate MTG dataset. To give an overview of the class conservation of 1-2-1 genes, this figure shows the average gene class conservation matrices among 10 pairs of species. See methods for calculation details. (a) Average specificity class conservation heatmap showing the percent overlap among one2one orthologs in various specificity class combinations (calculation process see methods). (b)-(c) Pie chart showing the aggregated percentage of cell type conservation and class conservation for genes with (b) cell type level specificity or (c) group level specificity. (d) Average distribution class conservation heatmap (calculation process see methods). (e)-(g) Pie chart showing the aggregated percentage of cell type conservation of genes stayed expressed in (e) over 90% of cell types; genes stayed expressed in (f) a single cell type and genes expressed in (g) some cell types. (h) Species tree of the 5 species analyzed in this figure.

For genes enriched in a specific cell type or group, an average of 47% or 36% remained in the same category in the other species. The numbers were lower for cell type or group enhanced genes, with 30% or 17% maintaining their specificity category, respectively. In contrast, genes with low specificity had a remarkable degree of consistency (averagely 84% overlap between species). Therefore, genes with extreme cell type expression specificity, whether high or low, exhibited the most conserved specificity class between species.

We further examined whether genes retained specificity for the same cell type (Figure 3b, c, see Supplementary figure 3 for each species pair results). On average, for genes with cell type level specificity, slightly over 90% of enriched genes that remained enriched were found in the same cell type, while the percentage for enhanced genes decreased to around 60% (Figure 3b). The cell type changes for genes that were group enriched or enhanced were more pronounced, with around 28% or 40% of events occurring in a different cell type, respectively (Figure 3c). Moreover, almost half of the events in which the gene transitioned between group enriched and group enhanced also changed the corresponding cell types.

Inspection of the frequency of switching between cell type pairs suggested that most genes switch their specificity between cell types known to be transcriptomically similar, such as between cerebral cortex endothelial cell and vascular leptomeningeal cell (see Supplementary figure 6a for frequency, Supplementary figure 6b for examples). Although, the higher number of genes switching between non-neuronal cell types could be confounded by the fact that they had more specific genes than the neurons.

Gene Ontology (GO) enrichment analysis on the most frequently conserved cell type-specific genes within each broad cell type among all species elucidated their roles in the primate MTG (Supplementary figure 7). In general, genes involved in foundational neuronal activity, such as synaptic signalling (e.g. *HTR2C; HTR3A; HTR3B; SLC32A1; SV2B; SV2C; PVALB; SLC1A2; SLC1A3)*, neural and glial development (e.g. *CPNE4; CPNE7; KIT; PDGFRA; PDGFRB; NOTCH3)* and extracellular matrix and cytoskeleton function e.g. (*MYO5B; IQGAP2; GFAP; LAMP5; MBP; CNTN2)* mainly maintained their cell type specificity (see full list in Supplementary table 1). Notably, there were a few transcription factors (TFs), which were known to be crucial in regulating neural development, found to be conserved between the 5 species. This includes *SP8* and *SP9* ^44^ and *EBF1/2/3* ^45^ which regulates interneuron differentiation and migration; *RUNX1/2/3* in regulating sensory neuron development ^46^; *ZIC1/2* regulates neurogenesis and differentiation ^47^, etc. We have also found crucial TFs for non-neuronal cell types, such as *KLF*2/4/6/8 in epithelial cells, and *TBX15/18 i*n vascular leptomeningeal cells. The results suggest that the basic mechanisms of neural signalling and development across mammalian cells are highly conserved at expression level. Furthermore, the enriched terms are highly consistent with a recent study of the ancestral neural-specific gene modules using bulk RNA-seq data across vertebrates and insects (Figure 2i by ^48^).

In terms of expression distribution, we saw that widely expressed genes maintained their broad expression while selectively expressed genes exhibited divergence. Figure 3d illustrates the averaged gene distribution class conservation between 1-2-1 orthologs in pairs of species (see Supplementary figure 8 for each species pair results). Regarding cell type expression maintenance, for genes expressed in over 90% of cell types, the frequency of expression in a different cell type was only 1% among all expression cases (Figure 3e). Yet as high as 67.9% of genes changed cell types between species if they were expressed in a single cell type (Figure 3f). Genes expressed in less than 30% and 30%-90% of cell types demonstrated more frequent cell type switching, particularly when they also changed between these distribution classes (Figure 3g). GO analysis revealed that genes consistently exhibiting low specificity and being expressed in over 90% of cell types between species were significantly enriched in terms associated with fundamental, constitutive cellular processes. These include transcription; RNA splicing; translation; post-translational modification; protein metabolism; cytoskeleton organization; cytosolic transport and mitochondrial function (Supplementary Figure 9). Such genes are characteristic of housekeeping roles, and their stability across species is unsurprising.

In summary, our analyses highlighted substantial divergence in the gene expression specificity and distribution profiles of 1-2-1 orthologs across species. The degree of class conservation and cell type conservation between species highly vary among genes of different classes. Genes with low expression specificity and broad expression distribution seemed to have the most conserved expression pattern between species, while highly cell type-specific genes and group-selectively expressed genes had relatively better conservation.

### Remote species show weaker 1-2-1 ortholog class conservation

To understand the impact of species divergence time on gene class conservation, we applied the gene classification analysis to 2 additional datasets. The 3 datasets provided perspectives for different species divergence times: ∼33 million years ago (MYA) for the primate MTG data ^49^, ∼450 MYA for the zebrafish/xenopus embryogenesis data ^50^ and ∼600-700 MYA for the Cnidarian data ^51^.

Comparing results from the 3 datasets demonstrated that the divergence time between species had a large impact on the one-to-one ortholog specificity and distribution class conservation patterns. As expected, in comparison to what we found in the data from evolutionally close primates, the zebrafish/xenopus pair showed overall less specificity class conservation, and a larger percentage of genes which switched cell type in different specificity categories (Supplementary figure 4). Among Cnidarian species, we observed even fewer 1-2-1 orthologs stayed in the same specificity class. In particular, more orthologs had a drastic change of their specificity classes, such as between cell type enriched and low cell type specificity (Supplementary figure 5). For the zebrafish/xenopus and Cnidarian species, the distribution classes were also generally less conserved, and cell type switching was more frequent. Compared with specificity classes, it seems that 1-2-1 orthologs are less likely to undergo extreme class switches between distribution classes. To conclude, the divergence of 1-2-1 ortholog expression is already prominent among five primates and is stronger for evolutionarily distant species.

### Gene class conservation patterns are stable respective to cell type granularity and can inform dataset comparability

By varying resolutions, one can classify cells differently, which adds an extra layer of complexity for cross-species comparison. Since gene classification by expression specificity and distribution depends on somewhat arbitrary chosen thresholds and cell annotations, genes can end up in different classes depending on the choices and the particular datasets. We were interested to what extent the gene class conservation results will be affected. Here, we varied the cell type classification to show that gene class conservation analysis gives stable trends across annotation granularities, as long as it is comparable between the respective species. Gene class conservation can further inform the comparability of populations and granularity between different species datasets.

As described previously, we found that in the primate MTG data, most cell-type-enriched genes were found in non-neuronal cells, which was also observed in the original publication as non-neuronal cells had 10-folds more marker genes ^7^. This could be explained by the fact that neuron classification might be higher in granularity than that of non-neuronal cells, although such classification is the most reasonable biologically. Therefore, we first used a reduced granularity of the primate MTG dataset to obtain a gene class conservation matrix and compared it with the original results from higher-granularity annotation. After using a more coarse classification of neuronal cell types, we found that non-neuronal cells still showed a higher number of specific genes. Importantly, the gene class conservation specificity and distribution matrix remained largely consistent (Supplementary figure 10 c, e). Moreover, a comparable percentage of specific genes switched cell type (Supplementary figure 10). These results demonstrate that the gene classification and conservation analysis is generally stable with respect to the cell type granularity, when the granularity used is comparable among species datasets, supporting the observation of frequent gene class switching of 1-2-1 orthologs between species.

The primate MTG dataset is unique in the sense that it has the same cell populations and matched annotation among the 5 species due to conscious sampling and data processing^52^. When the analysis concerns independently annotated data from different species, it is challenging to determine the true comparability in terms of cell population and annotation granularity. We found that the gene specificity class conservation matrix can provide information for assessing such comparability. Using GeneSpectra, we generated gene class conservation matrices from independently annotated human and mouse bone marrow datasets ^53^. Three additional cell types were included in the mouse dataset but were not profiled in the human dataset, a fact clear in this case but potentially uncertain in new contexts. When considering all cell types, the gene specificity class conservation matrices displayed a stripe of genes with specificity in the mouse but not in the human dataset (Supplementary figure 11a). In contrast, when only shared cell types were used, the matrices exhibited a clear diagonal pattern (Supplementary figure 11b). We conclude that the stripe pattern of genes with specificity on the gene specificity class conservation matrices can inform us that one species data has additional population. Following a similar intuition, if granularity is clearly higher in one species data, the gene class specificity and distribution conservation matrices will show a triangle pattern by having more specific genes, or more widely expressed genes in that species, respectively.

### Gene classes explain species effect on cross-species transcriptomic space

It is known that concatenated scRNA-seq data from different species exhibit strong global expression shift ^6^ known as species effects ^4^: cells show higher expression similarity within species, rather than within matching cell types, on the shared transcriptomic space built with 1-2-1 orthologs. Previously, we compared different computational strategies to remove this species effect and found it challenging to distinguish the source of such divergence ^4^. Here, we demonstrate that gene classification results can explain the species effect, which is one step further towards disentangling technical and biological expression divergence across-species.

For each pair of species, we obtained a list of 1-2-1 orthologs that stayed specific in the same specificity class (i.e. cell type or group enriched or enhanced, referred to as *conserved specific* genes hereafter). If the gene was also specific to the exact same cell type or groups, it was considered to be *strictly conserved specific*. We reason that the expression of these genes represent the evolutionarily conserved cell type transcriptomic profiles between species. Conversely, we also obtained a list of 1-2-1 orthologs that switched from specific to non-specific or lowly expressed in the other species (referred to as *diverged to unspecific* genes hereafter). We infer that these genes mainly contribute to the global expression divergence between the two species. Using the full list of 1-2-1 orthologs, strictly conserved specific genes, conserved specific genes and diverged to unspecific genes, we applied dimensional reduction to the single-cell expression profile of data from each pair of species, and measured the strength of the species effect.

Figure 4a shows an example of the UMAP visualisation of data from human and chimpanzee (other species pair results available in Supplementary figure 12). It is clear that species effects present on the full set of 1-2-1 orthologs data. When using conserved specific genes, cell types such as Sst; Pavalb; Vip neurons and Oligo; OPC; Astro and Chandelier showed partial overlap on the expression space, suggesting a much reduced species effect. On the other hand, when using diverged to unspecific genes, we observed a stronger species effect which led to the two species data being linearly separable on the first 2 dimensions on the PC space (Figure 4b). It appeared that the global species effect was extracted by restricting the features to diverge to unspecific genes. The data using only strictly conserved specific genes was highly similar with the one with conserved specific genes both in UMAP space and in PC space, possibly due to similar highly variable features being selected for dimension reduction. We quantified the degree of species effect by using PC regression score ^54^ (Figure 4c) and observed agreeing trends quantitatively with the dimensional reduction analysis.

**Figure 4.**
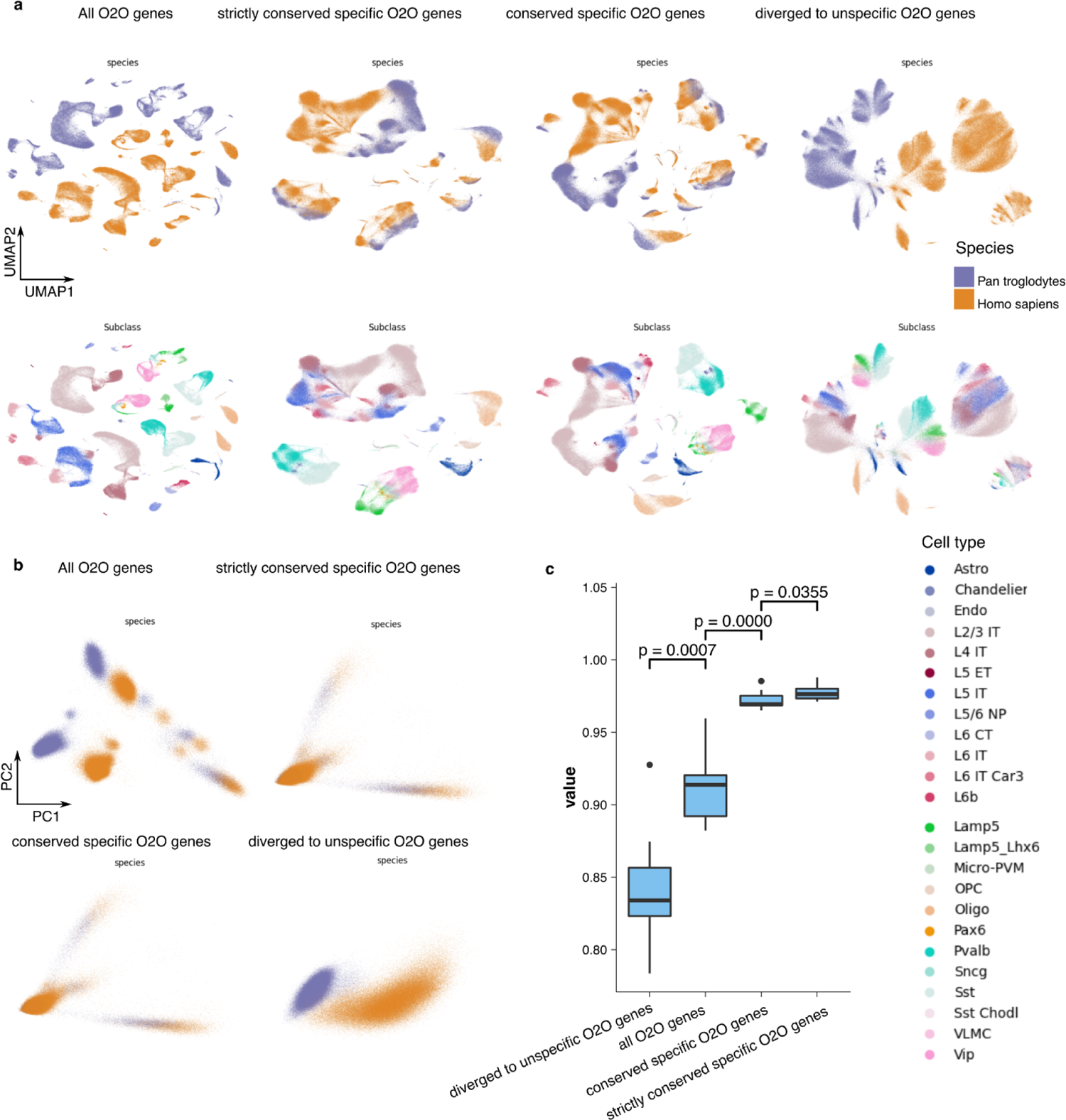
Gene class switching explains species effect on transcriptomic space. (a) UMAP visualisation using different sets of 1-2-1 orthologs for the human and chimpanzee data. Showing the strong species effect using all genes, the reduced species effect using (strictly) conserved specific genes, as well as a purified specie effect shown in diverged to unspecific genes (b) The first 2 PCs coloured by species using different sets of 1-2-1 orthologs for the human and chimpanzee data. (c) PCR score for data with different sets of genes. The bar in the boxplot shows the arithmetic mean, lower and upper hinges correspond to the first and third quartiles, whiskers extend from the hinge to the largest value no further than 1.5 * interquartile range and outliers beyond this range are plotted as individual data points. Wilcoxon signed-rank test was used to determine statistical significance (p value shown). 1-2-1, one-to-one; UMAP, Uniform Manifold Approximation and Projection; PC, principal component.

In conclusion, species effect on cross-species joint scRNA-seq data can largely be explained by ortholog gene switching from specific to non-specific expression. Although, the change of expression level of class-conserved genes can provide additional explanation. It is worth noting that it is not possible to fully separate the technical effects and the biological effects that drive species separation on joint transcriptomic space. However, by controlling for experimental batch effects the gene classification results can maximally capture true biological expression shifts.

### Positive correlation between expression specificity similarity and peptide product sequence conservation

Leveraging the gene classification results, we investigated the relationship between sequence conservation and the expression specificity similarity across cell types among different types of orthologs, with respect to the evolutionary proximity between species using the primate MTG dataset.

To measure peptide sequence conservation among orthologs, we computed the gene peptide product sequence alignment using BLASTp ^55^. Specificity patterns among orthologs were represented as vectors, where each value corresponds to the specificity score of the gene in a particular cell type (see Supplementary figure 13 for a schematic). The specificity score is defined as the ratio of the gene’s expression level in enriched or enhanced cell type or group to the enrichment or enhancement threshold (details in Methods and code repository). Intuitively, this vector characterises the gene by which cell types it is expressed and how specifically. If this vector includes all cell types of the organism (at the given level of granularity), then in a sense it can be viewed as a concise characterisation of the function of this gene in the organism. By measuring the similarity between these vectors, specifically, using the cosine function, we quantified the expression specificity similarity between genes from different species. We plotted the cosine similarity of expression specificity against the bit score for cell type specific orthologs for each pair of species in the primate MTG data (Figure 5 for cell type enriched genes, see Supplementary figure 14 for cell type enhanced genes).

**Figure 5.**
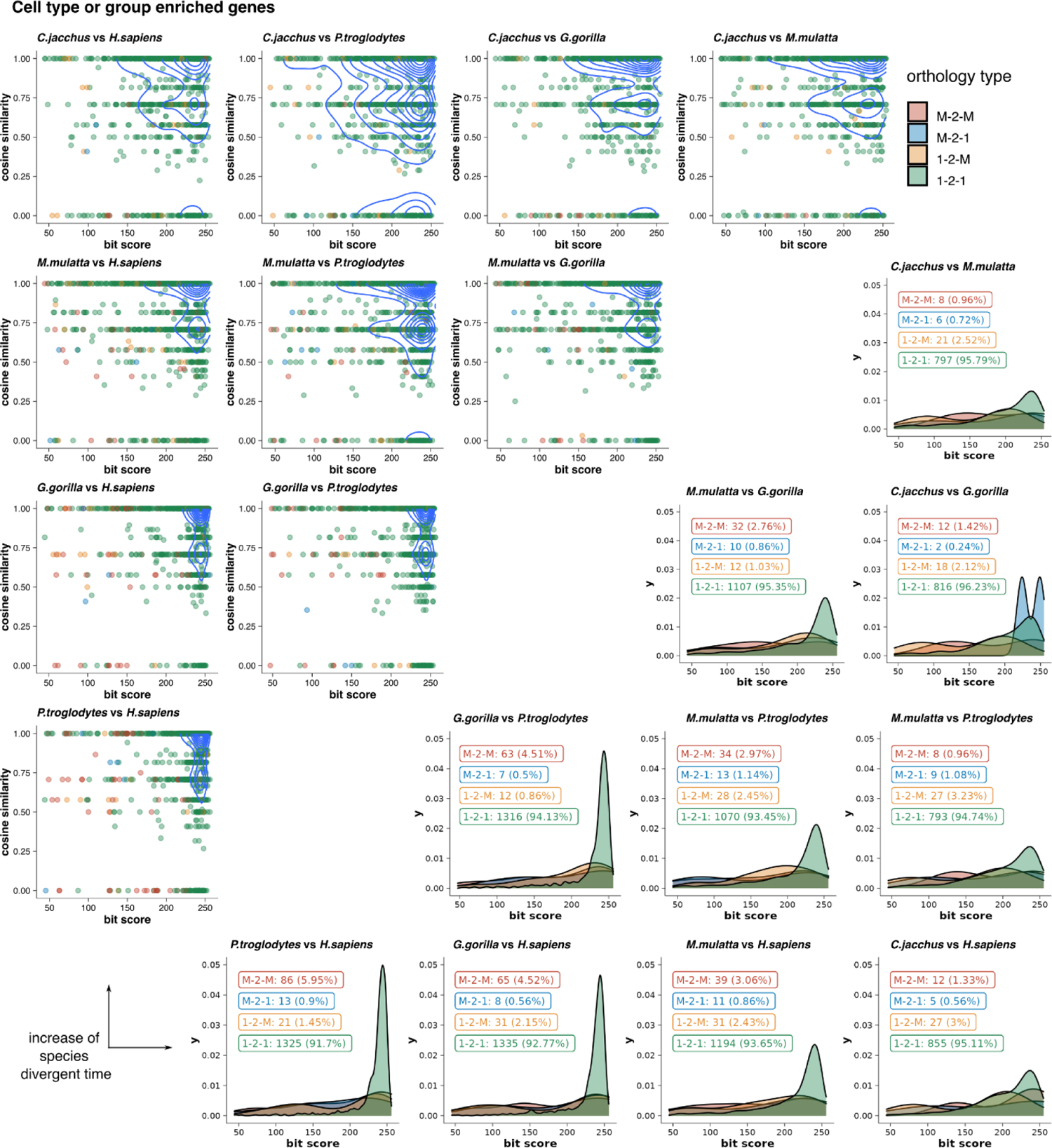
Comparing sequence conservation with cell type expression specificity similarity across species distance for enriched genes. The top left triangle plots the sequence conservation (bit score) against the expression similarity (cosine similarity) for each pair of cell type enriched orthologous in each pair of species. The bottom right triangle shows the distribution of bit scores for different types of orthologs for the same set of genes. 1-2-1: one-to-one, 1-2-M: one-to-many, M-2-1: many-to-one, M-2-M: many-to-many.

We identified a major group of genes with high sequence conservation and conserved cell type specificity, located in the top-right corner of each panel within the top-left triangle of Figure 5. Meanwhile, a significant group of genes displayed moderate sequence conservation but no expression similarity, positioned along the lower edge. GO enrichment analysis indicated that genes with high expression similarity and high sequence conservation were mainly involved in gliogenesis and neurogenesis; cell-cell adhesion; vascular and blood-brain barrier transport and immune response, all of which closely relate to the major functions of the cell types involved (Supplementary figure 15). There was no clear enrichment for genes with high sequence similarity but no expression specificity.

There are relatively few genes in the middle area. Interestingly, with the increase of species divergence time, we observed a spread out pattern of genes towards having a medium sequence conservation and expression specificity similarity, as well as an increase of the percentage of genes with high sequence conservation but zero expression specificity similarity. This suggests that distant species have a more diverged genome both in terms of sequence and gene expression. Further supporting these observations, applying the same analysis to the embryogenesis data revealed that most genes were found in the middle region with a more dispersed distribution, suggesting a weaker association between expression specificity similarity and peptide sequence conservation (Supplementary figure 16).

Considering different types of orthologs, we found that non-1-2-1 orthologs in general have lower sequence conservation than 1-2-1 orthologs (bottom right triangle in Figure 5), but we saw no clear bias in expression specificity similarity. Many non-1-2-1 orthologs with a range of sequence conservation showed high expression conservation. Our results agree with the observations by Stamboulian, M. et al. ^19^ that the types of homologs are largely irrelevant to function prediction tasks, highlighting the importance of including non-1-2-1 orthologs into functional analysis.

In a nutshell, our results suggested an overall positive relationship between sequence conservation and expression similarity, particularly between more closely related species. Such positive correlation becomes weaker with the increase of evolutionary distance, and for less specifically-expressed genes. Nevertheless, a sizeable group of genes did not follow this pattern, and non-1-2-1 orthologs didn’t show differences of expression specificity similarity compared with 1-2-1 orthologs, despite lower sequence conservation.

## Discussion

In this work, we evaluated the ortholog conjecture in the context of scRNA-seq data between species, considering cell type expression specificity and distribution as the functional readouts of genes. We found that orthologous genes with low cell type specificity tend to retain these expression patterns across species, consistent with their likely roles in essential housekeeping functions. Likewise, strongly cell type or group-specific genes also exhibited relatively high level of expression pattern conservation, albeit to a lesser extent and less so in distantly related species. In contrast, genes with intermediate specificity exhibited the greatest variability in specificity patterns between species. Regarding expression distribution, genes that were broadly expressed in one species tend to maintain this broad distribution across species, whereas genes with highly exclusive expression frequently changed the cell type in which they were expressed. On balance, these findings suggest that the degree to which ortholog conjecture applies depends on the cell type expression specificity and distribution of the respective orthologs, highlighting the nuanced and context-dependent nature of gene expression evolution.

Our results provide new insights to the ortholog conjecture debate, as previous studies ^20,21,23,56–58^ did not yet have scRNA-seq data available. It is possible that not distinguishing genes with different cell type specificities led to contradicting observations between these studies using bulk data. From the perspective of gene regulation, this highlights a key distinction: genes with tightly regulated, cell type-specific expression differ in evolution patterns from those with broad, consistent expression across cell types. Furthermore, with the increase of species evolutionary distance, ortholog conjecture generally becomes less applicable for all gene classes. This is reflected by an overall decrease of genes stayed in the same class, and somewhat surprisingly, more drastic gene class changes between 1-2-1 orthologs. Taken together, these findings underscore the need to evaluate and apply the ortholog conjecture separately for genes with distinct cell type expression patterns, while considering species relationship as an integral part of the ortholog conjecture discussion ^25^.

Understanding the expression specificity and distribution conservation of orthologs highlights key avenues for advancing computational methods in cross-species cell type evolutionary mapping. Evident specificity class switching between orthologs indicated that full-transcriptome ortholog expression correlation between cell types is a poor approximation of their evolutionary relatedness. Instead, one should seek evidence based on the evolutionary definition of cell types^1^. The definition emphasises the distinction between the regulatory mechanism that establishes cell type identity, and gene modules that produce specific cell type phenotypes in a species ^1^. The theory proposes that terminal selector TFs ^31,59^ and their effector genes maintain their cell type-specific expression between species and is indicative of evolutionary cell type homology. Other genes that contribute to cellular function but are not deterministic of cell identity can have more variable expression.

Expression signals from both aspects are detected simultaneously in scRNA-seq data, and our analysis to obtain *strictly conserved cell type-specific* genes essentially disentangles the two. Strictly conserved cell type-specific orthologs can serve as candidates of terminal selectors (if they are TFs) and effector genes for developing methods for aligning evolutionarily related cell types across species, a process that can be further refined with additional data modalities for direct gene regulation information. Such a theoretical framework can also be applied to group-specific genes as identity-defining regulatory mechanisms operate on the level of cell type families ^60^. This perspective also helps explain why we observed a comparable degree of ortholog specificity conservation when analysing data at a lower level of cell type granularity. Here, the group-specific genes essentially became classified as “cell type-specific”. At a higher level, we also note that using gene expression as a proxy for gene function has its limitations: it is effective for genes responsible for maintaining cell identity, but less so for genes whose expression is mostly driven by environmental stimuli and adaptive demands.

Focusing on the evolutionary mapping of cells based on a few strictly conserved specific genes can also help resolve the challenge of circularity faced by the field. The issue of circularity arises when cell types are defined via aligning the full scRNA-seq atlas between species, and then these cell type classifications are used to study gene expression conservation cross-species. Therefore, it is important to only use the gene expression signals that evolutionarily define cell type identity to match cell types between species, then the comparison at the full transcriptome level to detect species-specific expression will be more robust. With non-1-2-1 orthologs showing a comparable expression specificity similarity with 1-2-1 orthologs, non-1-2-1 orthologs should also be included into the functional mapping of cell types between species. This draws parallel from the conclusion that including different types of orthologs greatly improves gene functional prediction ^19^. As the genome and its context evolve, new genes specific to a species can significantly contribute to the identity-defining regulatory mechanisms of species-specific cell types. It is thus essential to also consider the contribution of species-specific genes into calculating the expression similarity of cell types.

Important questions related to the ortholog conjecture are yet to be addressed in scRNA-seq data. First is to further examine the expression pattern conservation among non-1-2-1 orthologs, and compare the conservation between orthologs with that of in-paralogs in scRNA-seq data. It is currently technically challenging to accurately assign scRNA-seq reads among in-paralogs that have high sequence similarity within one species. In this work, we were underpowered to address this with publicly available datasets, despite that the computational framework remains applicable. More generally, for future cross-species comparison endeavours, ensuring dataset comparability is essential. It is important to generate datasets with future cross-study compatibility in mind. It is also challenging to find the correct granularity for cell type comparison that is biologically meaningful and technically feasible. We show that gene class conservation matrices can serve as an initial check for the comparability of cell populations and annotation granularity, yet biological knowledge about the species is crucial to interpret these data and perform cell type mapping between species. Additionally, it is worth noting that many cross-species expression differences detected computationally could be evolutionarily neutral ^7^. The addition of functional data will help to disentangle the signals that are relevant to species adaptation.

## Methods

### Single-cell RNA-seq data used in this study

All datasets analysed in this study are publicly available. Sn-RNA-seq count matrices from humans, chimpanzees, gorillas, rhesus macaques, and marmosets were downloaded from the publication release “Great_Ape_MTG_Analysis’’ from the NeMO archive ^61^. Author cell ontology annotations were obtained from Cellxgene of the same dataset ^62^. Only 10x V3 3’ data and cells passed the original QC of the published dataset were analysed in this study to minimise technical effects. InDrops data of zebrafish and xenopus embryos were obtained from the GEO database under accession code GSE11229434 and GSE11307435, respectively. Cnidarian datasets were downloaded from Mendeley data ^63^ uploaded by the authors. Human and mouse bone marrow data were obtained from ArrayExpress under accession number E-MTAB-8629 and E-MTAB-8630.

### Gene orthology mapping

The gene orthology mapping and orthology type information for the primates primate MTG data, the embryogenesis data and the bone marrow data were obtained from ENSEMBL (release 110) via the biomart (v0.9.2) python package. For the Cnidarian data, we used the orthology mapping results provided by the authors calculated with Broccoli ^43,63^. The gene identifiers in the Cnidarian scRNA-seq were not mapped to publically available genomes, so we could not extend the analysis beyond author-provided 1-2-1 orthologs.

### Constructing metacells

Metacells partition the raw single-cell data into disjoint, homogeneous groups ideally representing re-sampling from the same cellular state ^40^. We downloaded metacells (V0.9.4) from GitHub (https://github.com/tanaylab/metacells). Metacells were constructed with default hyperparameters following the tutorial https://tanaylab.github.io/metacells-vignettes/iterative.html. Lateral genes related to cell cycle and stress, and genes highly correlate with these genes were masked during metacell construction. Metacell cell type annotations were conveyed from single-cell annotations by maximum percentage per metacell. In the primate MTG data, approximately 15-25 single cells, nearly 100% of the same cell type, constituted one metacell (Supplementary figure 1).

### Cell type pooling as alternative to metacells

Due to the continuity of cell type expression in embryonic settings, we employed an alternative approach for sparsity reduction which is cell type pooling. We added the count values of cells from the same type as input for GeneSpectra, which was then subjected to depth normalisation. Such cell type pooling approaches could be used in datasets where metacell is not applicable.

### The GeneSpectra module for gene classification

The GeneSpectra module was developed under Python 3.8.16. It takes an AnnData (v0.9.1) object containing metacells and calculates the average normalised metacell expression for each group (in this context, cell type). After transforming the results into a long data format, it applies a classification function to each gene when using multithreading to speed up the computation. Finally, it outputs a dataframe containing specificity and distribution classification results. Various plotting and analysis functions were also implemented to aid comprehension and downstream analysis.

The gene expression specificity class is determined as follows: if the normalised expression level of the gene in a certain cell type, or the average of a group of cell types, was 4 folds higher than the maximum of all the other cell types, it was considered enriched. On the other hand, if the elevation was only 4 folds higher than the average of all the other cell types, it was considered enhanced. Cell type or group enriched or enhanced genes were collectively referred to as cell type specific genes in this paper. The gene expression distribution class was determined by the percentage of cell types where the normalised expression level was above 1 CPM. In both criteria, genes that had an expression < 1 CPM in all cell types are considered lowly expressed.

### Gene class conservation matrix

To calculate the gene specificity and distribution class conservation matrix for 1-2-1 orthologs between each species pair, we followed these steps: First, 1-2-1 orthologs were identified using ENSEMBL, and the number of orthologs in each class was counted for both species. Next, we determined the intersection of orthologs across all class combinations for the species pair. For each species, the intersection was expressed as a percentage of the total genes in that class. Finally, the heatmap shows the harmonic mean of these percentages for the two species. Using the harmonic mean, rather than raw gene counts, accounts for differences in the total number of 1-2-1 orthologs in different classes across species.

To show an average class conservation among several pairs of species in the primate MTG and Cnidarian datasets, the species were paired in a directional way: from species A to species B, with species B more closely related to humans. As such, the averaged heatmap can be comprehended as genes move from class in species A to species B along the direction of time.

### GO enrichment

GO enrichments were performed using the R package clusterProfiler (v4.10.0) with org.Hs.eg.db (3.18.0, GO data source date 2023-07-27), while visualization was supported by GOSemSim (v2.28.0), enrichplot (v1.23.1.992), and DOSE (v3.28.2). Human served as the reference species for all primate MTG analyses and only human vs other species were analysed. The GO_basic ontology was used. Background genes were restricted to those showing non-zero expression in the human dataset. P-values were corrected for multiple comparisons using the Benjamini-Hochberg method, and results were considered significant if P < 0.01.

### ScRNA-seq data analysis and visualisation

To perform analysis and visualisation of scRNA-seq data using different sets of 1-2-1 orthologs from pairs of species, we used Scanpy (v1.4.6) with the following parameters: sc.pp.highly_variable_genes(adata, min_mean = 0.0125, max_mean = 3, min_disp = 0.5); sc.pp.scale(adata, max_value = 10); sc.tl.pca(adata, svd_solver = ‘arpack’); sc.pp.neighbours(adata, n_neighbors = 15, n_pcs = 40); sc.tl.umap(adata, min_dist = 0.3, spread = 1).

### BLASTp

The following protein translation of ENSEMBL genes were downloaded from ENSEMBL: human (Homo_sapiens.GRCh38.pep.all.fa); chimpanzee(Pan_troglodytes.Pan_tro_3.0.pep.all.fa); gorilla (Gorilla_gorilla.gorGor4.pep.all.fa); rhesus macaque (Macaca_mulatta.Mmul_10.pep.all.fa); marmoset (Callithrix_jacchus.mCalJac1.pat.X.pep.all.fa); zebrafish (Danio_rerio.GRCz10.pep.all.fa); xenopus (Xenopus_tropicalis.Xenopus_tropicalis_v9.1.pep.all.fa). We ran BLASTp (V2.15.0) using default parameters and used the bit score as the sequence conservation score.

## Software availability

The GeneSpectra module is available at https://github.com/Papatheodorou-Group/GeneSpectra and the version of code used in this study is available via Zenodo with DOI: 10.5281/zenodo.13303986.

## Declarations

The authors declare no competing interests.

## Author contributions

All authors conceived this study with A.B. made the initial proposal. Y.S. developed the softwares, performed analysis and wrote the draft manuscript with supervision and inputs from I.P. and A.B., and insights from D.A.. All authors reviewed and edited the manuscript.

## Acknowledgements

This work was supported by: the European Molecular Biology Laboratory (Y.S., D.A., A.B., I.P.); the EMBL international PhD program (Y.S.); and the Biotechnology and Biological Sciences Research Council (BBSRC) grant ‘Fly Cell Atlas’ [BB/T014563/1] (I.P.).

