## Supplementary Figures for "Revising the ortholog conjecture in cross-species comparison of scRNA-seq data"

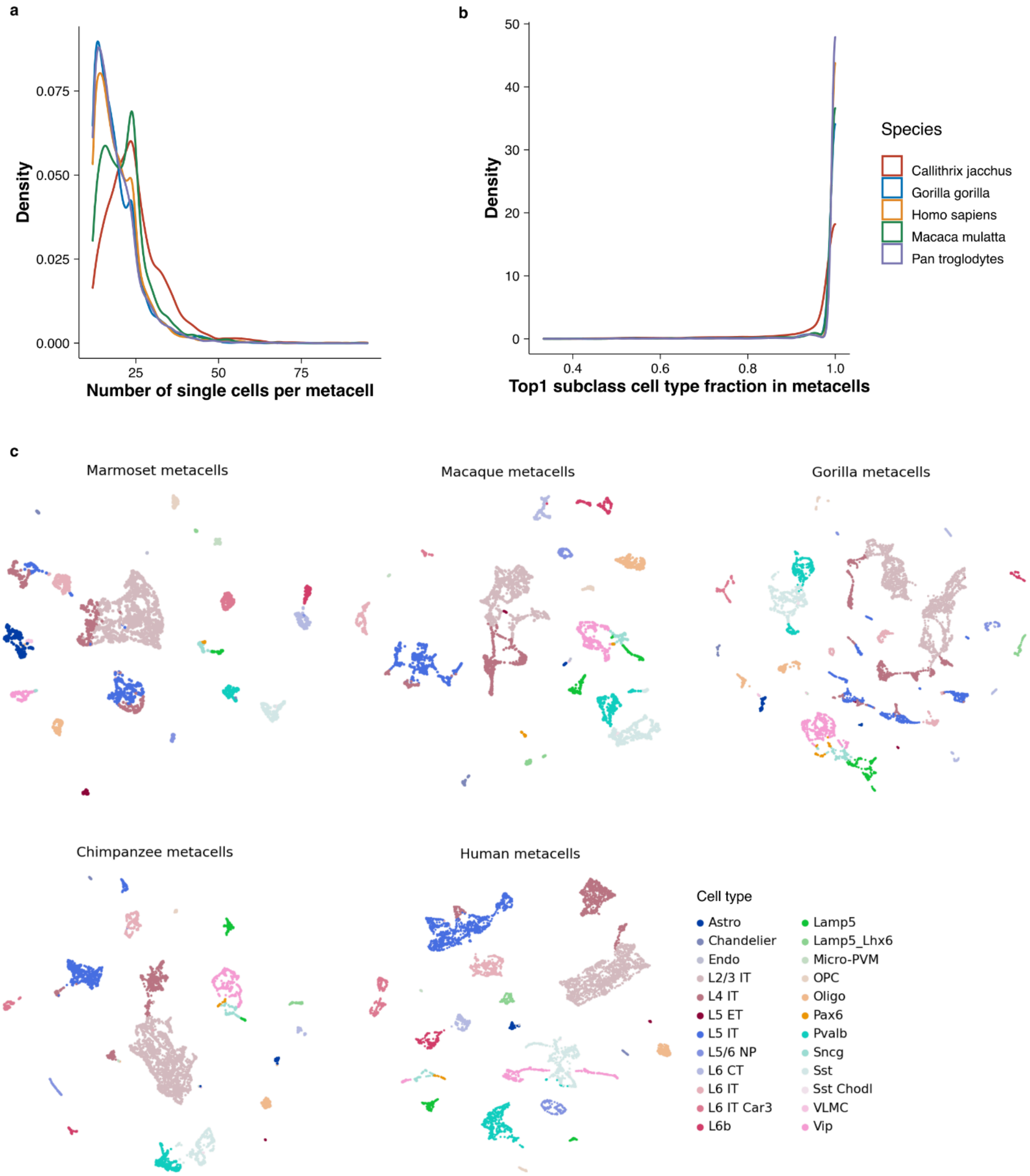


**Supplementary Figure 1 Metacells in the primate MTG data.** (a) Number of single cells per metacell in each species. (2) Fraction of the top 1 cell type in each metacell in each species. (c) UMAP visualisation of metacells in each species with cell type annotation (“Subclass" in original study). MTG, middle temporal gyrus.

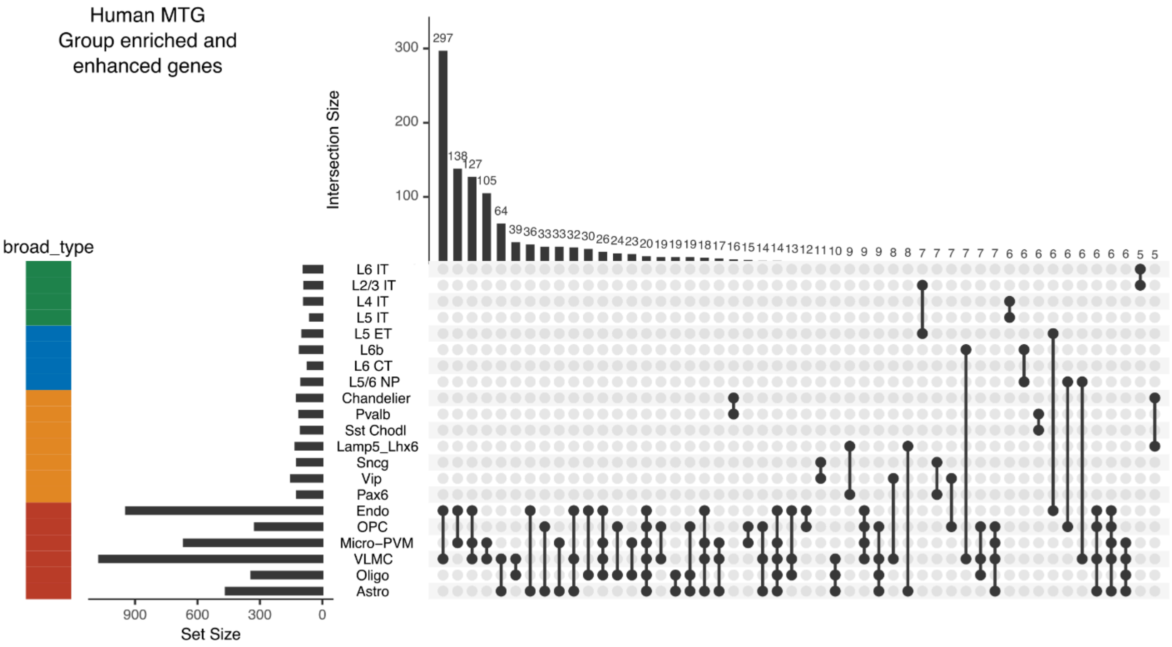
Human MTG
Group enriched and
enhanced genes
Intersection Size
broad_type
L6 IT
L2/3 IT
L4 IT
L5 IT
L5 ET
L6b
L6 CT
L5/6 NP
Chandelier
Pvalb
Sst Chodl
Lamp5_Lhx6
Sncg
Vip
Pax6
Endo
OPC
Micro−PVM
VLMC
Oligo
Astro
Set Size

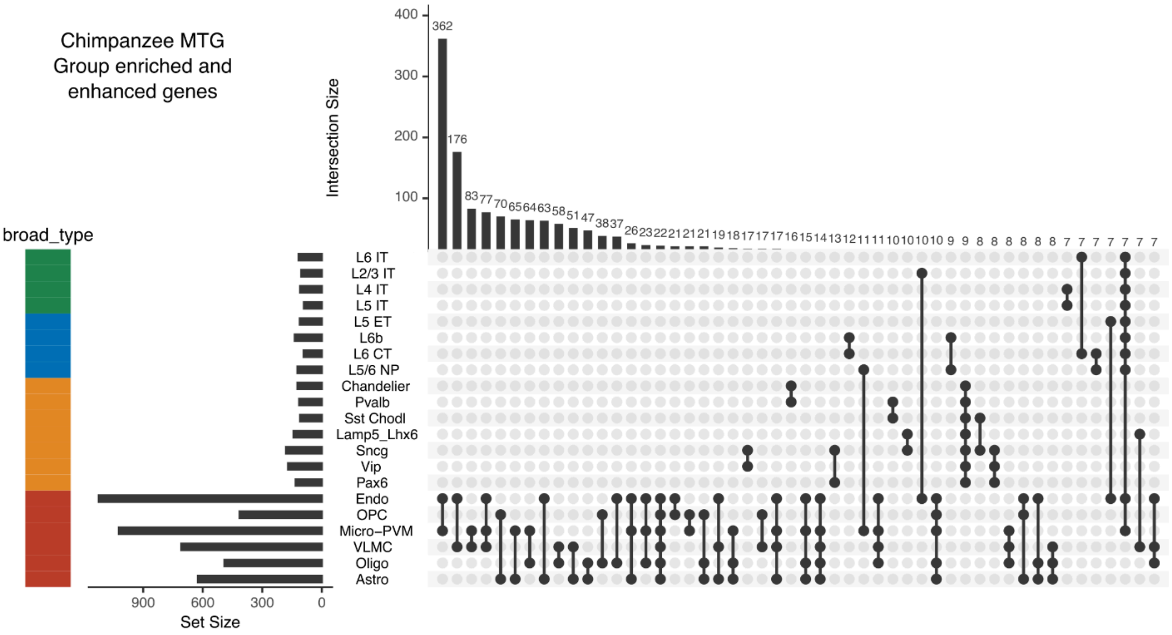
Chimpanzee MTG
Group enriched and
enhanced genes
Intersection Size
broad_type
L6 IT
L2/3 IT
L4 IT
L5 IT
L5 ET
L6b
L6 CT
L5/6 NP
Chandelier
Pvalb
Sst Chodl
Lamp5_Lhx6
Sncg
Vip
Pax6
Endo
OPC
Micro−PVM
VLMC
Oligo
Astro
Set Size

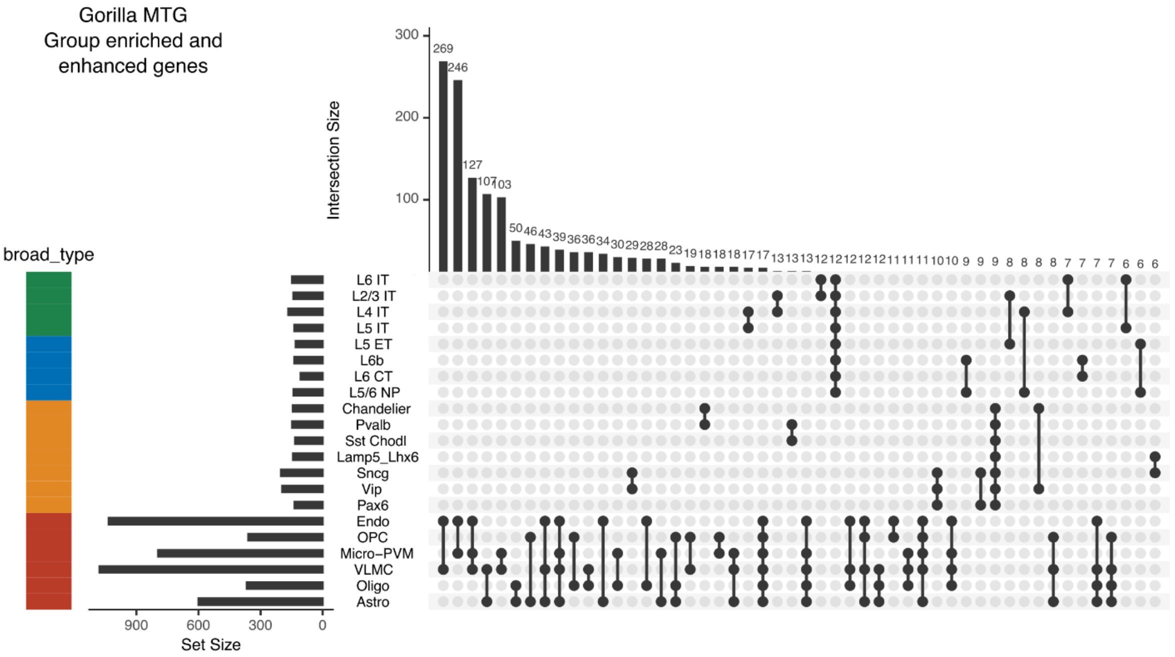
Gorilla MTG
Group enriched and
enhanced genes
Intersection Size
300
200
100
broad_type
L6 IT
L2/3 IT
L4 IT
L5 IT
L5 ET
L6b
L6 CT
L5/6 NP
Chandelier
Pvalb
Sst Chodl
Lamp5_Lhx6
Sncg
Vip
Pax6
Endo
OPC
Micro−PVM
VLMC
Oligo
Astro
900
600
300
0
Set Size

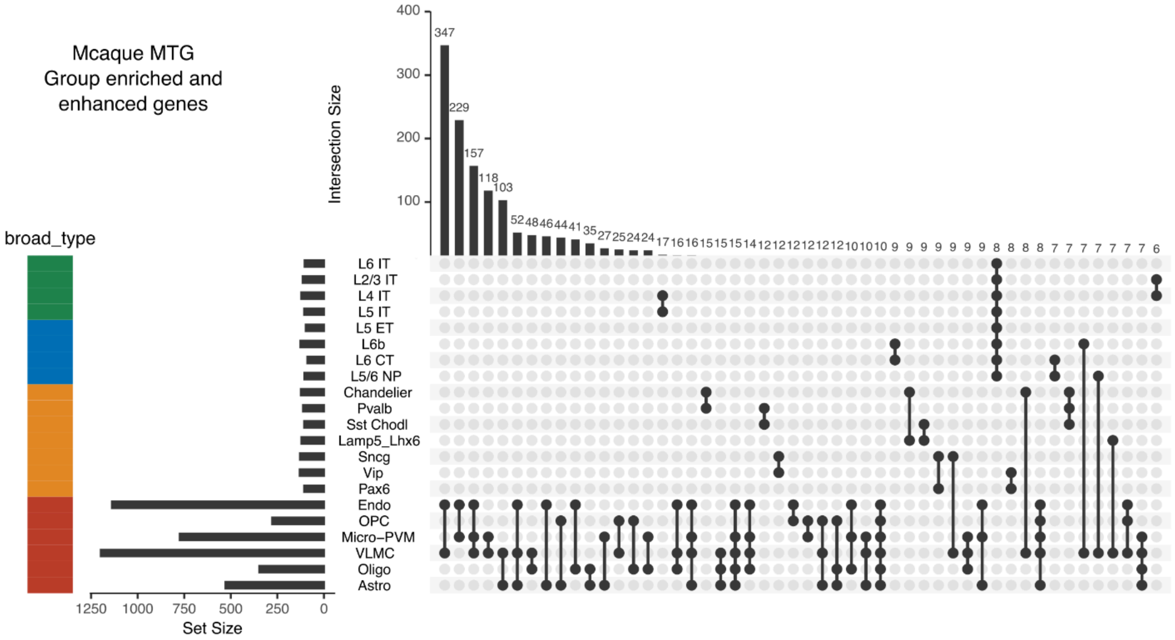
Mcaque MTG
Group enriched and
enhanced genes
Intersection Size
400
300
200
100
broad_type
L6 IT
L2/3 IT
L4 IT
L5 IT
L5 ET
L6b
L6 CT
L5/6 NP
Chandelier
Pvalb
Sst Chodl
Lamp5_Lhx6
Sncg
Vip
Pax6
Endo
OPC
Micro−PVM
VLMC
Oligo
Astro
1250
1000
750
500
250
0
Set Size

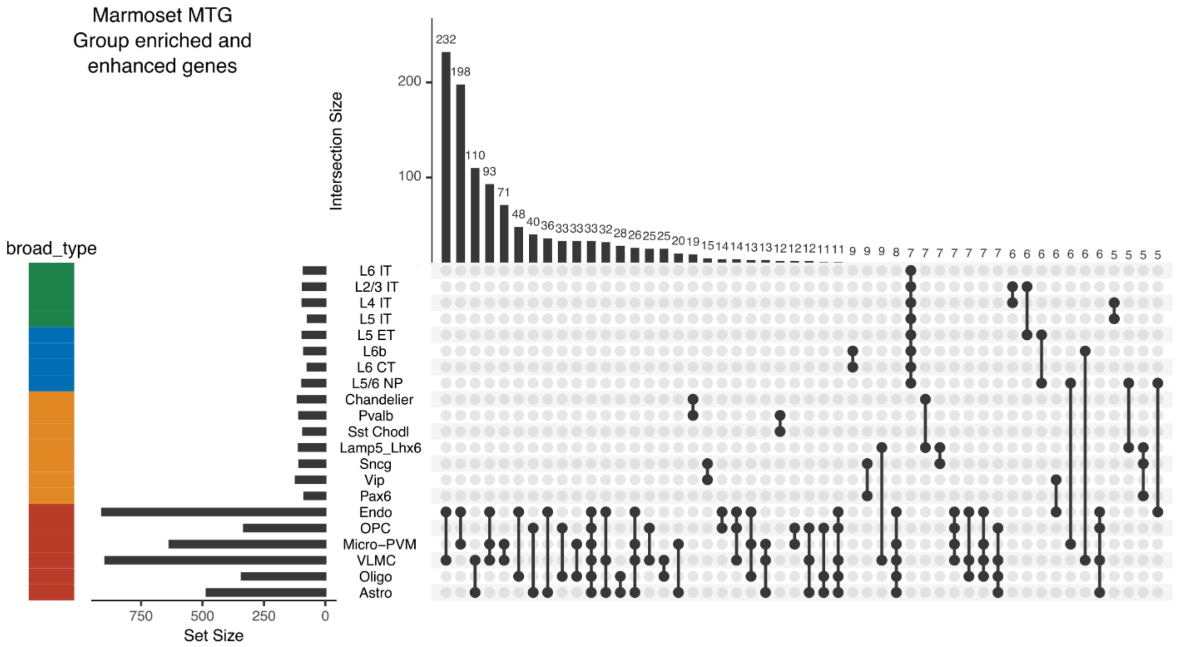


**Supplementary Figure 2 Group specific genes sharing among cell types.** Upset plots showing which cell types share the most group specific (enriched and enhanced) genes in each species. Intersections are ordered by intersection size while set size is ordered by broad type.

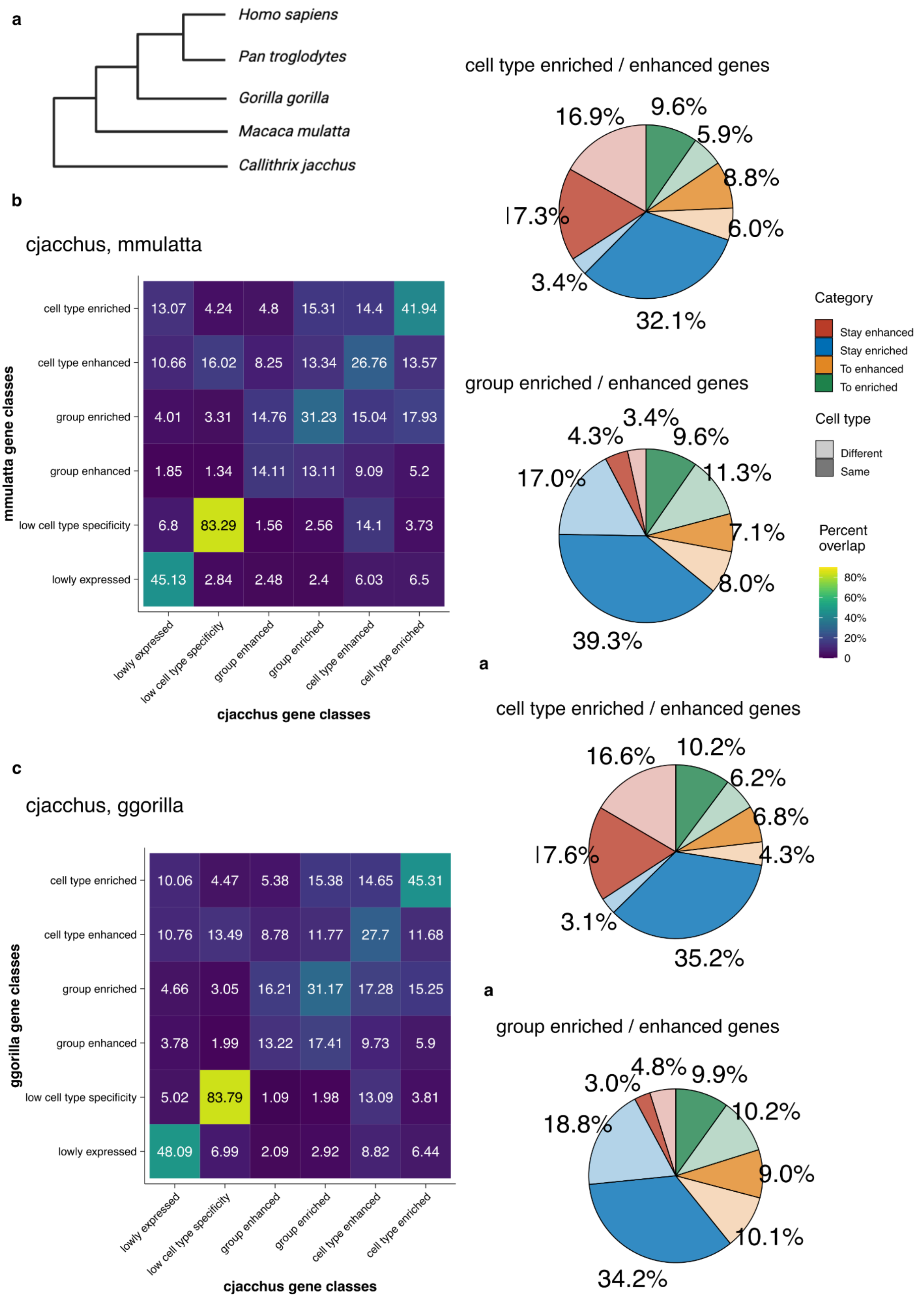
a
Homo sapiens
Pan troglodytes
Gorilla gorilla
Macaca mulatta
Callithrix jacchus
b
cjacchus, mmulatta
mmulatta gene classes
cell type enriched 13.07 4.24 4.8 15.31 14.4 41.94
cell type enhanced 10.66 16.02 8.25 13.34 26.76 13.57
group enriched 4.01 3.31 14.76 31.23 15.04 17.93
group enhanced 1.85 1.34 14.11 13.11 9.09 5.2
low cell type specificity 6.8 83.29 1.56 2.56 14.1 3.73
lowly expressed 45.13 2.84 2.48 2.4 6.03 6.5
lowly expressed
low cell type specificity
group enhanced
group enriched
cell type enhanced
cell type enriched
cjacchus gene classes
c
cjacchus, ggorilla
ggorilla gene classes
cell type enriched 10.06 4.47 5.38 15.38 14.65 45.31
cell type enhanced 10.76 13.49 8.78 11.77 27.7 11.68
group enriched 4.66 3.05 16.21 31.17 17.28 15.25
group enhanced 3.78 1.99 13.22 17.41 9.73 5.9
low cell type specificity 5.02 83.79 1.09 1.98 13.09 3.81
lowly expressed 48.09 6.99 2.09 2.92 8.82 6.44
lowly expressed
low cell type specificity
group enhanced
group enriched
cell type enhanced
cell type enriched
cjacchus gene classes
cell type enriched / enhanced genes
16.9%
9.6%
5.9%
8.8%
6.0%
17.3%
3.4%
32.1%
group enriched / enhanced genes
4.3%
3.4%
9.6%
11.3%
17.0%
7.1%
8.0%
39.3%
Category
Stay enhanced
Stay enriched
To enhanced
To enriched
Cell type
Different
Same
Percent overlap
80%
60%
40%
20%
0
a
cell type enriched / enhanced genes
16.6%
10.2%
6.2%
6.8%
17.6%
4.3%
3.1%
35.2%
a
group enriched / enhanced genes
3.0%
4.8%
9.9%
10.2%
18.8%
9.0%
10.1%
34.2%

**d**

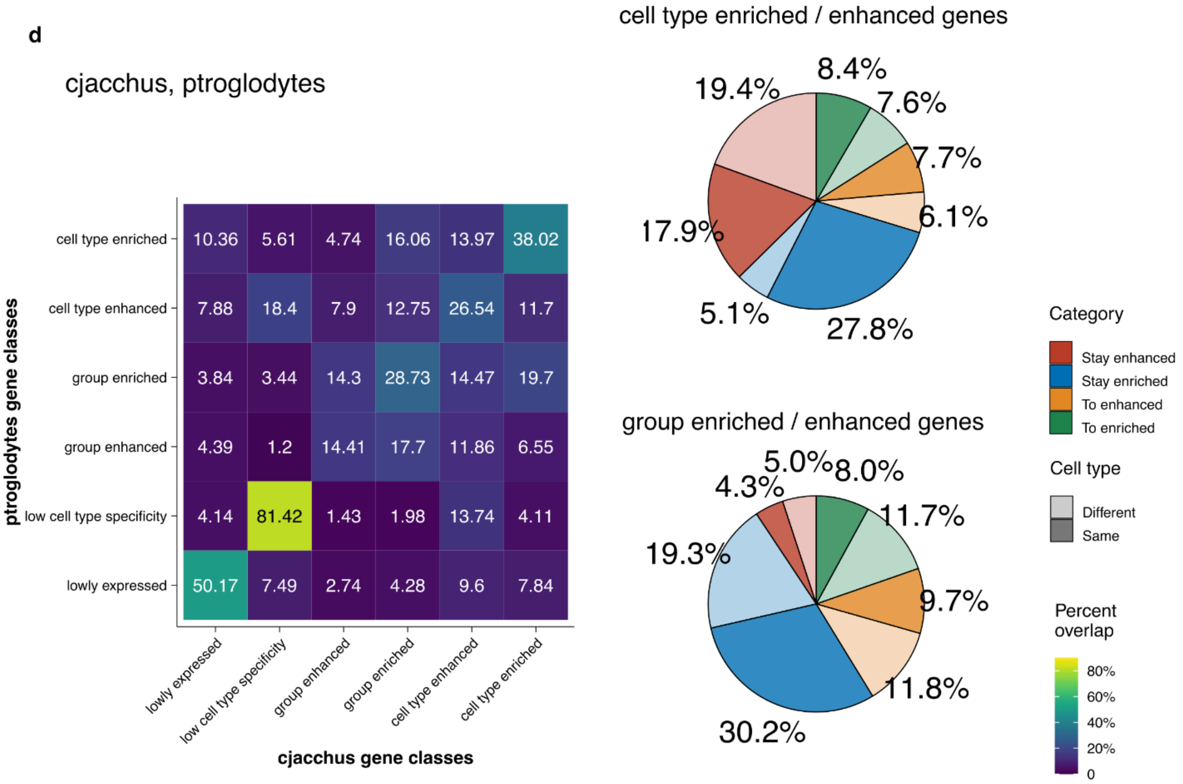
cjacchus, ptroglodytes
ptroglodytes gene classes
cell type enriched
cell type enhanced
group enriched
group enhanced
low cell type specificity
lowly expressed
10.36 5.61 4.74 16.06 13.97 38.02
7.88 18.4 7.9 12.75 26.54 11.7
3.84 3.44 14.3 28.73 14.47 19.7
4.39 1.2 14.41 17.7 11.86 6.55
4.14 81.42 1.43 1.98 13.74 4.11
50.17 7.49 2.74 4.28 9.6 7.84
lowly expressed
low cell type specificity
group enhanced
group enriched
cell type enhanced
cell type enriched
cjacchus gene classes
cell type enriched / enhanced genes
8.4%
7.6%
7.7%
6.1%
27.8%
5.1%
17.9%
19.4%
group enriched / enhanced genes
5.0%
8.0%
11.7%
9.7%
11.8%
30.2%
19.3%
4.3%
Category
Stay enhanced
Stay enriched
To enhanced
To enriched
Cell type
Different
Same
Percent overlap
80%
60%
40%
20%
0

**e**

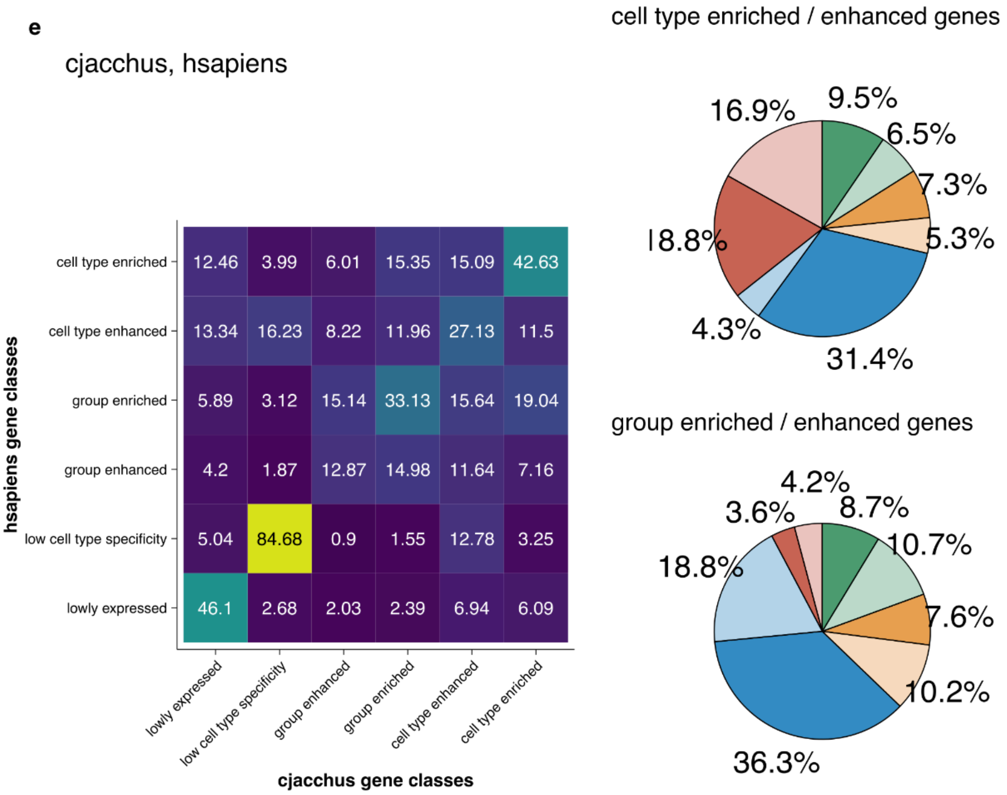
cjacchus, hsapiens
hsapiens gene classes
cell type enriched
cell type enhanced
group enriched
group enhanced
low cell type specificity
lowly expressed
12.46 3.99 6.01 15.35 15.09 42.63
13.34 16.23 8.22 11.96 27.13 11.5
5.89 3.12 15.14 33.13 15.64 19.04
4.2 1.87 12.87 14.98 11.64 7.16
5.04 84.68 0.9 1.55 12.78 3.25
46.1 2.68 2.03 2.39 6.94 6.09
lowly expressed
low cell type specificity
group enhanced
group enriched
cell type enhanced
cell type enriched
cjacchus gene classes
cell type enriched / enhanced genes
9.5%
6.5%
7.3%
5.3%
31.4%
4.3%
18.8%
16.9%
group enriched / enhanced genes
4.2%
8.7%
10.7%
7.6%
10.2%
36.3%
18.8%
3.6%

f

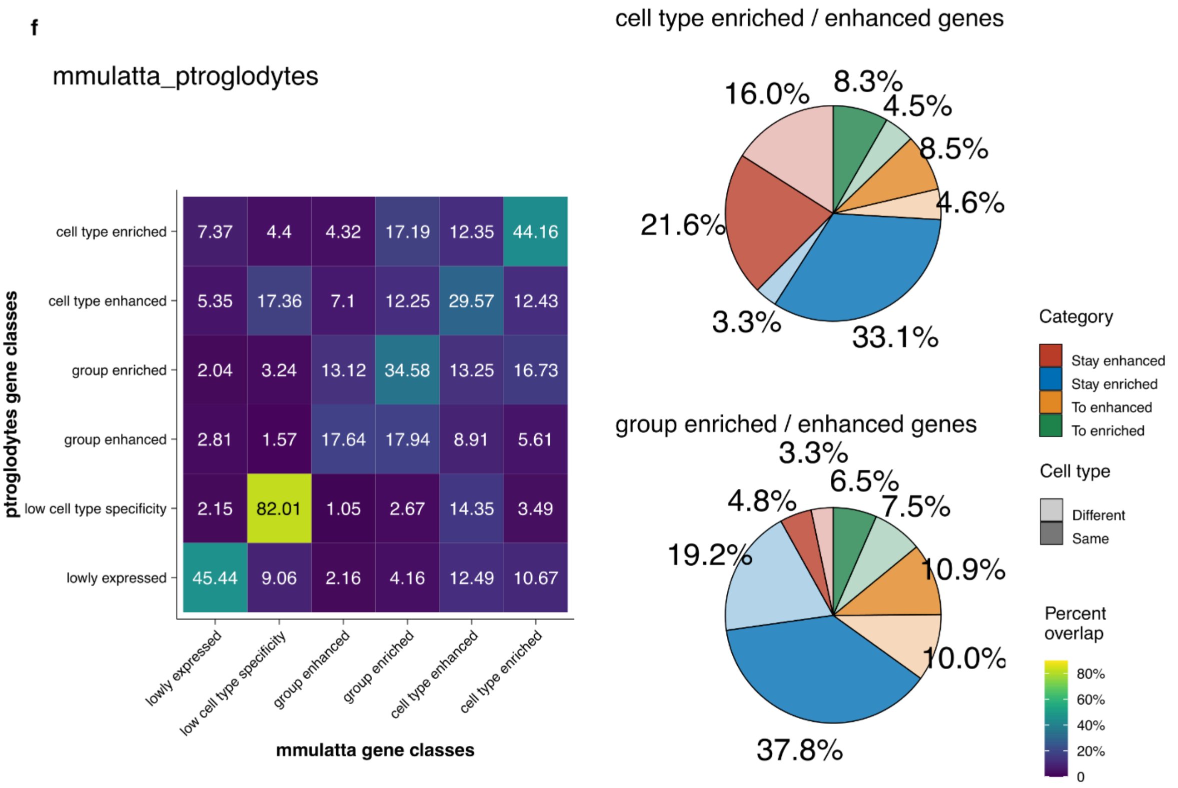

mmulatta_ptroglodytes
ptroglodytes gene classes
cell type enriched 7.37 4.4 4.32 17.19 12.35 44.16
cell type enhanced 5.35 17.36 7.1 12.25 29.57 12.43
group enriched 2.04 3.24 13.12 34.58 13.25 16.73
group enhanced 2.81 1.57 17.64 17.94 8.91 5.61
low cell type specificity 2.15 82.01 1.05 2.67 14.35 3.49
lowly expressed 45.44 9.06 2.16 4.16 12.49 10.67
lowly expressed
low cell type specificity
group enhanced
group enriched
cell type enhanced
cell type enriched
mmulatta gene classes
cell type enriched / enhanced genes
16.0%
8.3%
4.5%
8.5%
4.6%
21.6%
3.3%
33.1%
group enriched / enhanced genes
3.3%
6.5%
4.8%
7.5%
19.2%
10.9%
10.0%
37.8%
Category
Stay enhanced
Stay enriched
To enhanced
To enriched
Cell type
Different
Same
Percent overlap
80%
60%
40%
20%
0


g

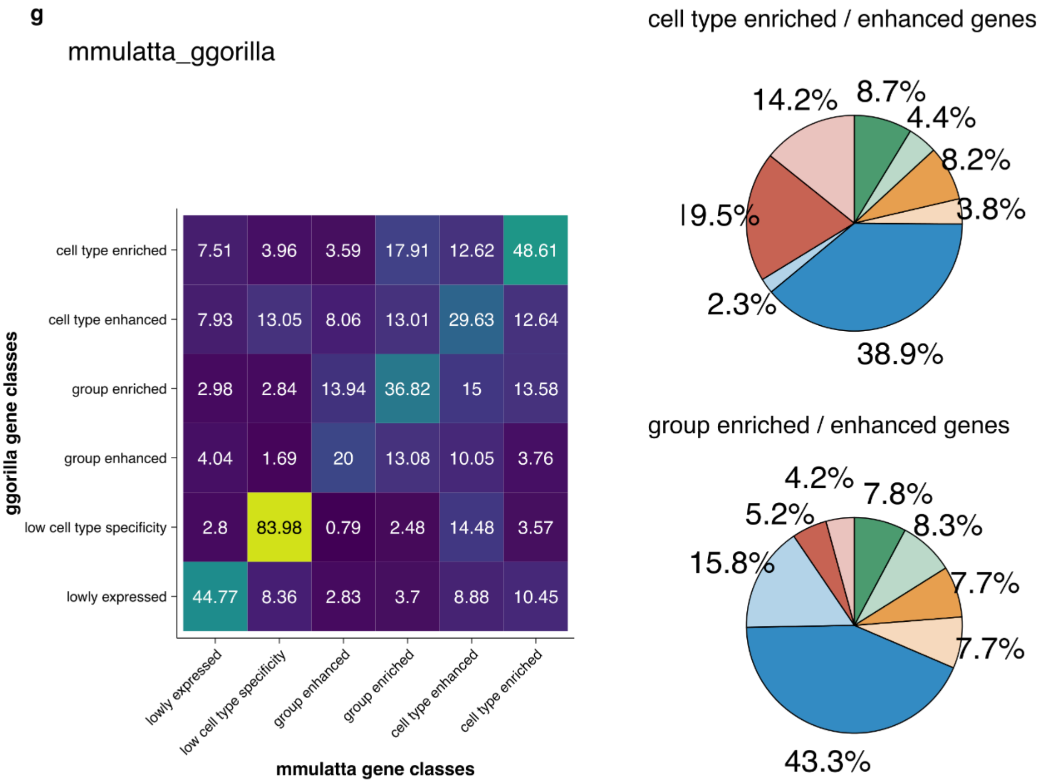

mmulatta_ggorilla
ggorilla gene classes
cell type enriched 7.51 3.96 3.59 17.91 12.62 48.61
cell type enhanced 7.93 13.05 8.06 13.01 29.63 12.64
group enriched 2.98 2.84 13.94 36.82 15 13.58
group enhanced 4.04 1.69 20 13.08 10.05 3.76
low cell type specificity 2.8 83.98 0.79 2.48 14.48 3.57
lowly expressed 44.77 8.36 2.83 3.7 8.88 10.45
lowly expressed
low cell type specificity
group enhanced
group enriched
cell type enhanced
cell type enriched
mmulatta gene classes
cell type enriched / enhanced genes
14.2%
8.7%
4.4%
8.2%
19.5%
3.8%
2.3%
38.9%
group enriched / enhanced genes
4.2%
7.8%
5.2%
8.3%
15.8%
7.7%
7.7%
43.3%

h

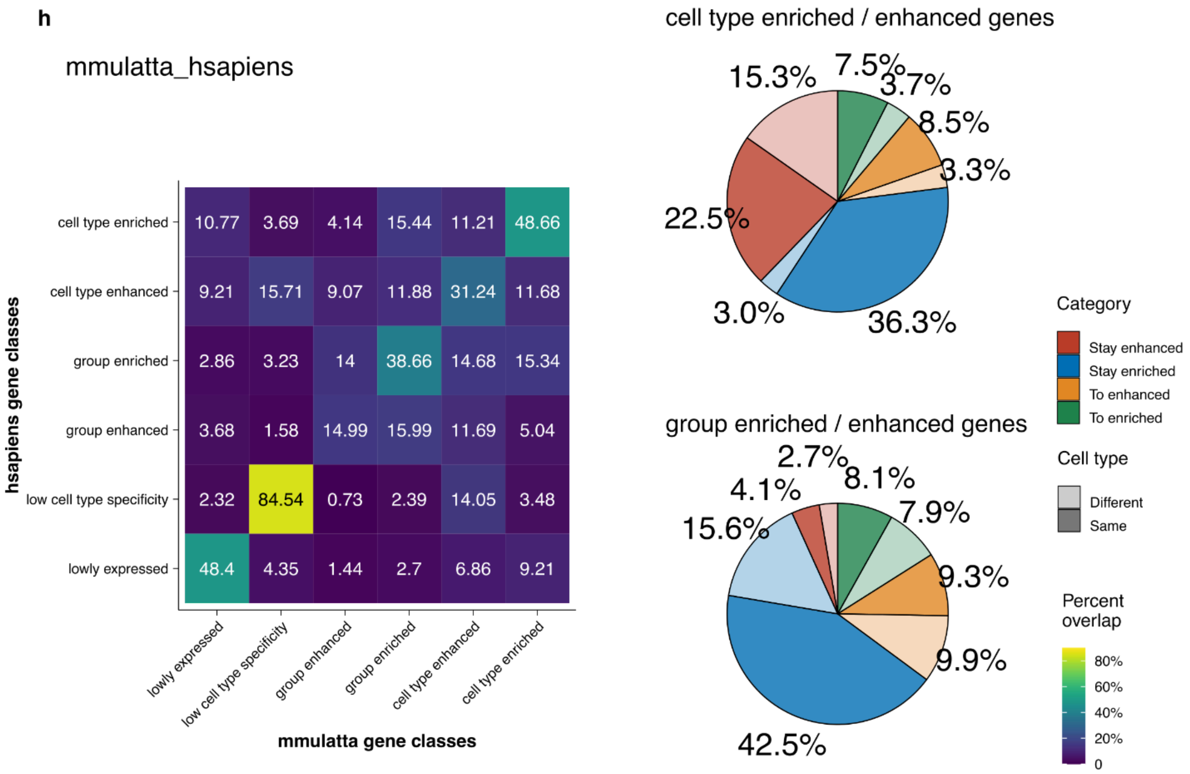
mmulatta_hsapiens
hsapiens gene classes
mmulatta gene classes
cell type enriched 10.77 3.69 4.14 15.44 11.21 48.66
cell type enhanced 9.21 15.71 9.07 11.88 31.24 11.68
group enriched 2.86 3.23 14 38.66 14.68 15.34
group enhanced 3.68 1.58 14.99 15.99 11.69 5.04
low cell type specificity 2.32 84.54 0.73 2.39 14.05 3.48
lowly expressed 48.4 4.35 1.44 2.7 6.86 9.21
lowly expressed
low cell type specificity
group enhanced
group enriched
cell type enhanced
cell type enriched
cell type enriched / enhanced genes
15.3%
7.5%
3.7%
8.5%
3.3%
22.5%
3.0%
36.3%
group enriched / enhanced genes
2.7%
8.1%
4.1%
7.9%
15.6%
9.3%
9.9%
42.5%
Category
Stay enhanced
Stay enriched
To enhanced
To enriched
Cell type
Different
Same
Percent overlap
80%
60%
40%
20%
0

i

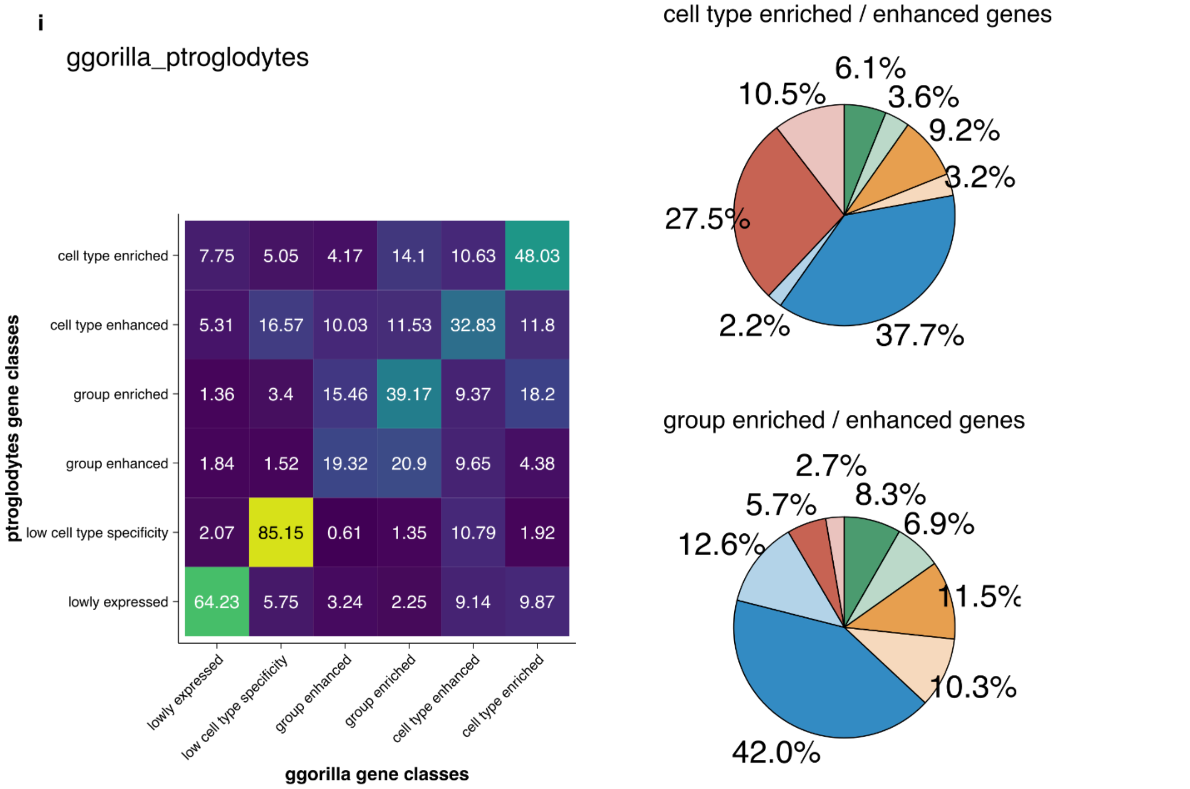
ggorilla_ptroglodytes
ptroglodytes gene classes
ggorilla gene classes
cell type enriched 7.75 5.05 4.17 14.1 10.63 48.03
cell type enhanced 5.31 16.57 10.03 11.53 32.83 11.8
group enriched 1.36 3.4 15.46 39.17 9.37 18.2
group enhanced 1.84 1.52 19.32 20.9 9.65 4.38
low cell type specificity 2.07 85.15 0.61 1.35 10.79 1.92
lowly expressed 64.23 5.75 3.24 2.25 9.14 9.87
lowly expressed
low cell type specificity
group enhanced
group enriched
cell type enhanced
cell type enriched
cell type enriched / enhanced genes
6.1%
10.5%
3.6%
9.2%
3.2%
27.5%
2.2%
37.7%
group enriched / enhanced genes
2.7%
5.7%
8.3%
6.9%
12.6%
11.5%
10.3%
42.0%

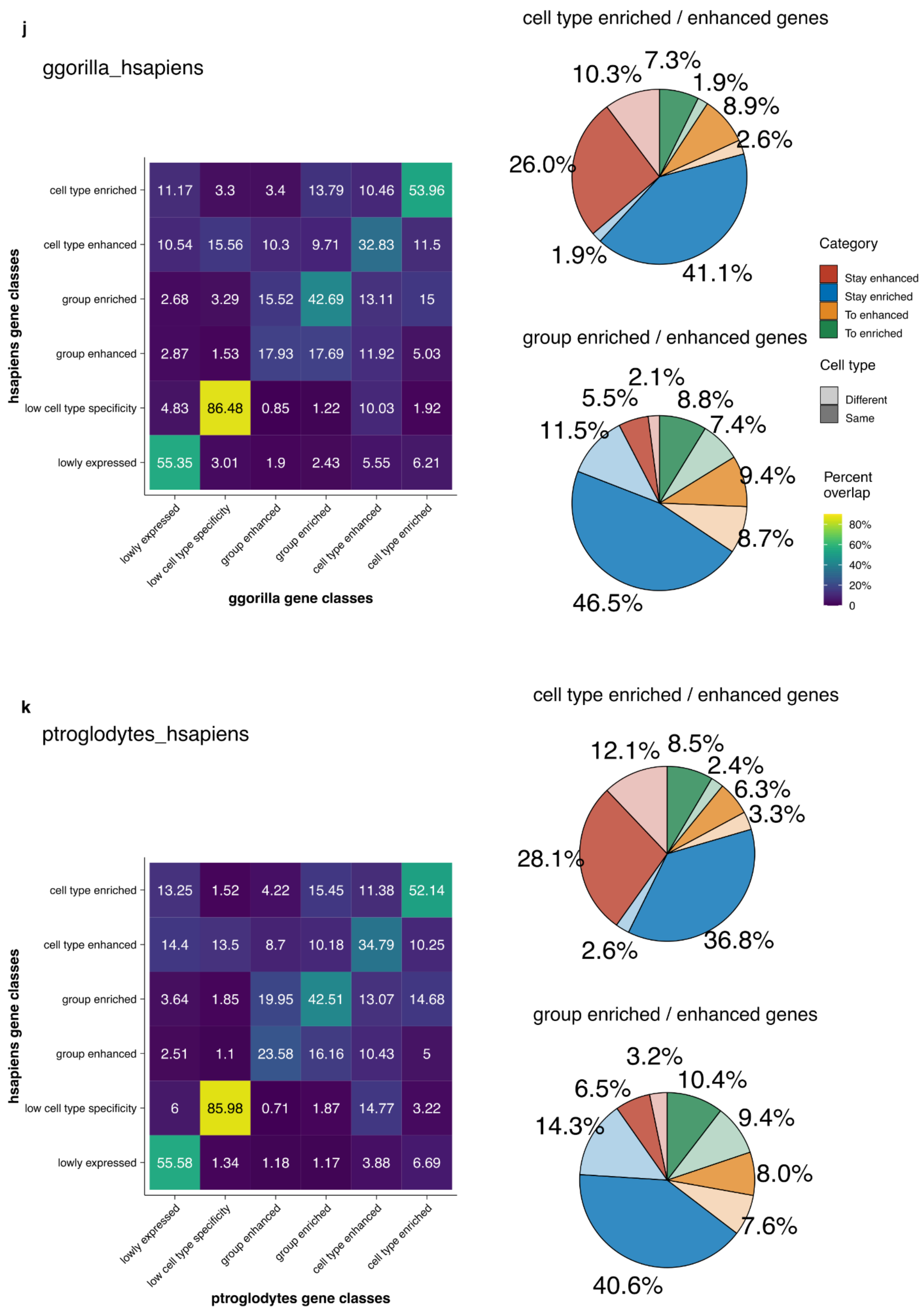


**Supplementary Figure 3 Gene specificity class conservation among O2O orthologs in primate MTG dataset, for each species pair.** Average specificity class conservation heatmap showing the percent overlap among one2one orthologs in various specificity class combinations (calculation process see methods). Pie chart showing the aggregated percentage of cell type conservation and class conservation for genes with cell type level specificity or group level specificity.

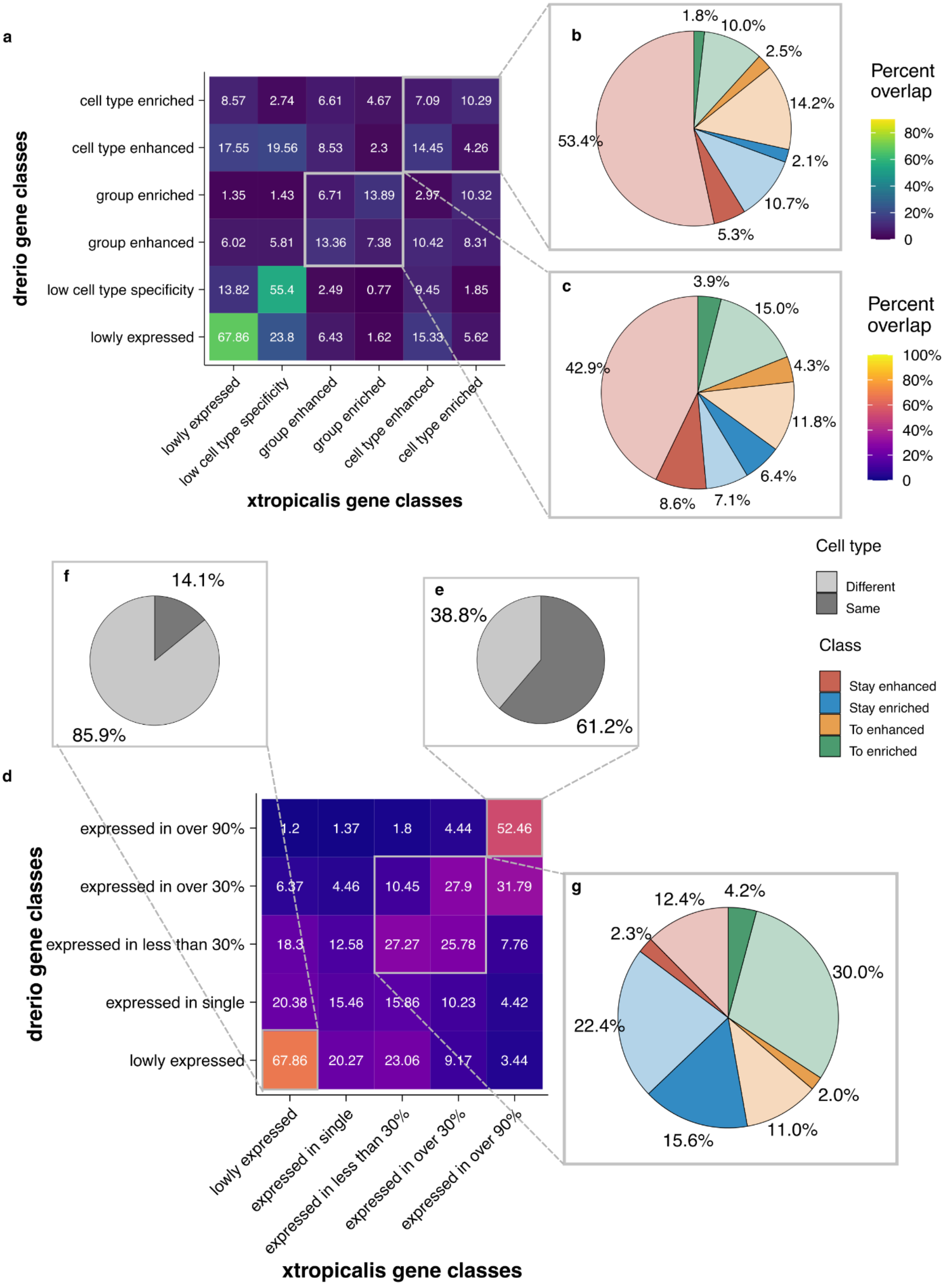


**Supplementary figure 4 Gene class conservation among O2O orthologs in embryogenesis dataset.** (a) Average specificity class conservation heatmap showing the percent overlap among one2one orthologs in various specificity class combinations (calculation process see methods). (b)-(c) Pie chart showing the aggregated percentage of cell type conservation and class conservation for genes with (b) cell type level specificity or (c) group level specificity. (d) Average distribution class conservation heatmap (calculation process see methods). (e)-(g) Pie chart showing the aggregated percentage of cell type conservation of genes stayed expressed in (e) over 90% of cell types; genes stayed expressed in (f) a single cell type and genes expressed in (g) some cell types.

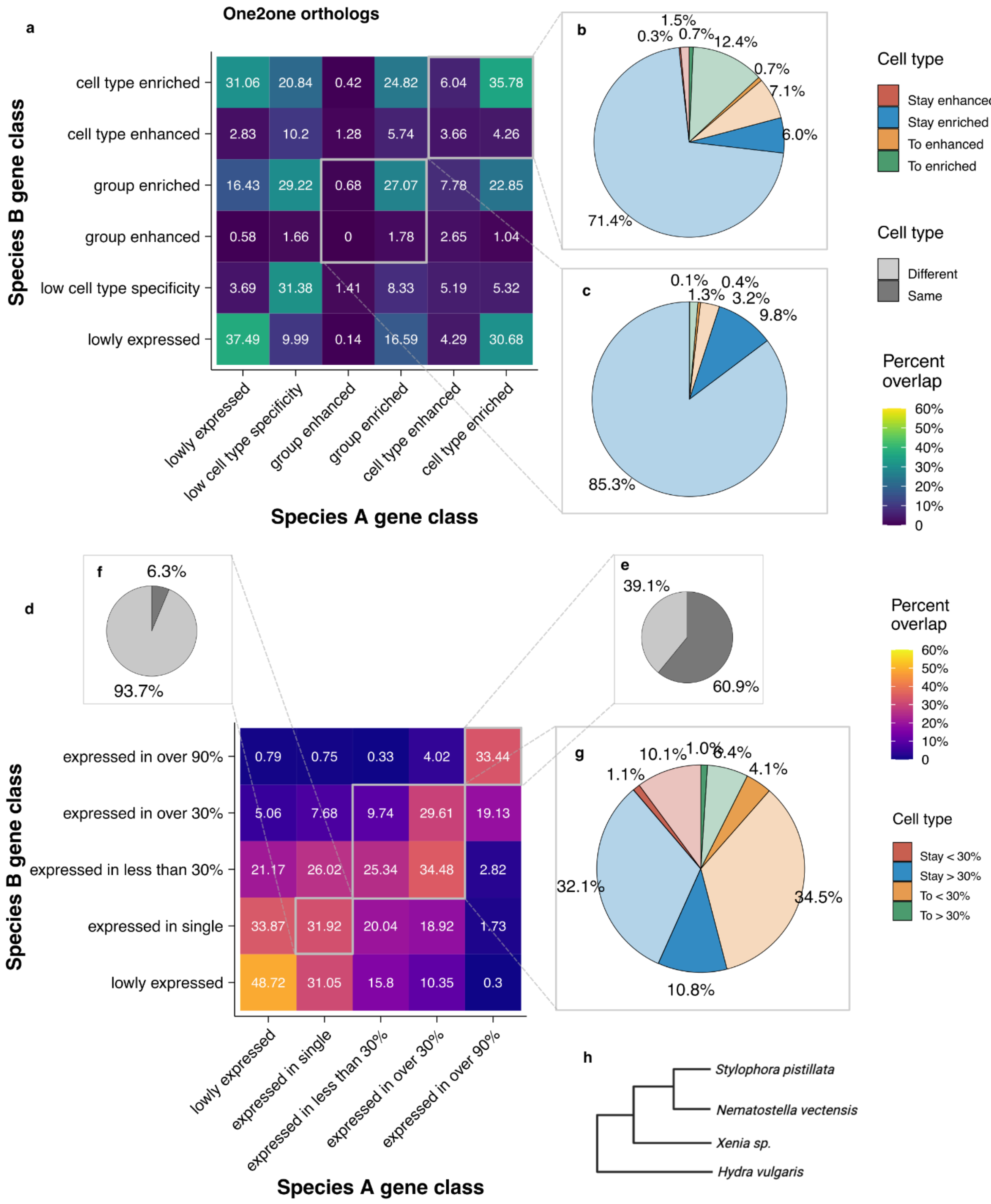


**Supplementary figure 5 Gene class conservation among O2O orthologs in Cnidarian dataset.** (a) Average specificity class conservation heatmap showing the percent overlap among one2one orthologs in various specificity class combinations (calculation process see methods). (b)-(c) Pie chart showing the aggregated percentage of cell type conservation and class conservation for genes with (b) cell type level specificity or (c) group level specificity. (d) Average distribution class conservation heatmap (calculation process see methods). (e)-(g) Pie chart showing the aggregated percentage of cell type conservation of genes stayed expressed in (e) over 90% of cell types; genes stayed expressed in (f) a single cell type and genes expressed in (g) some cell types. (h) species tree of this dataset.

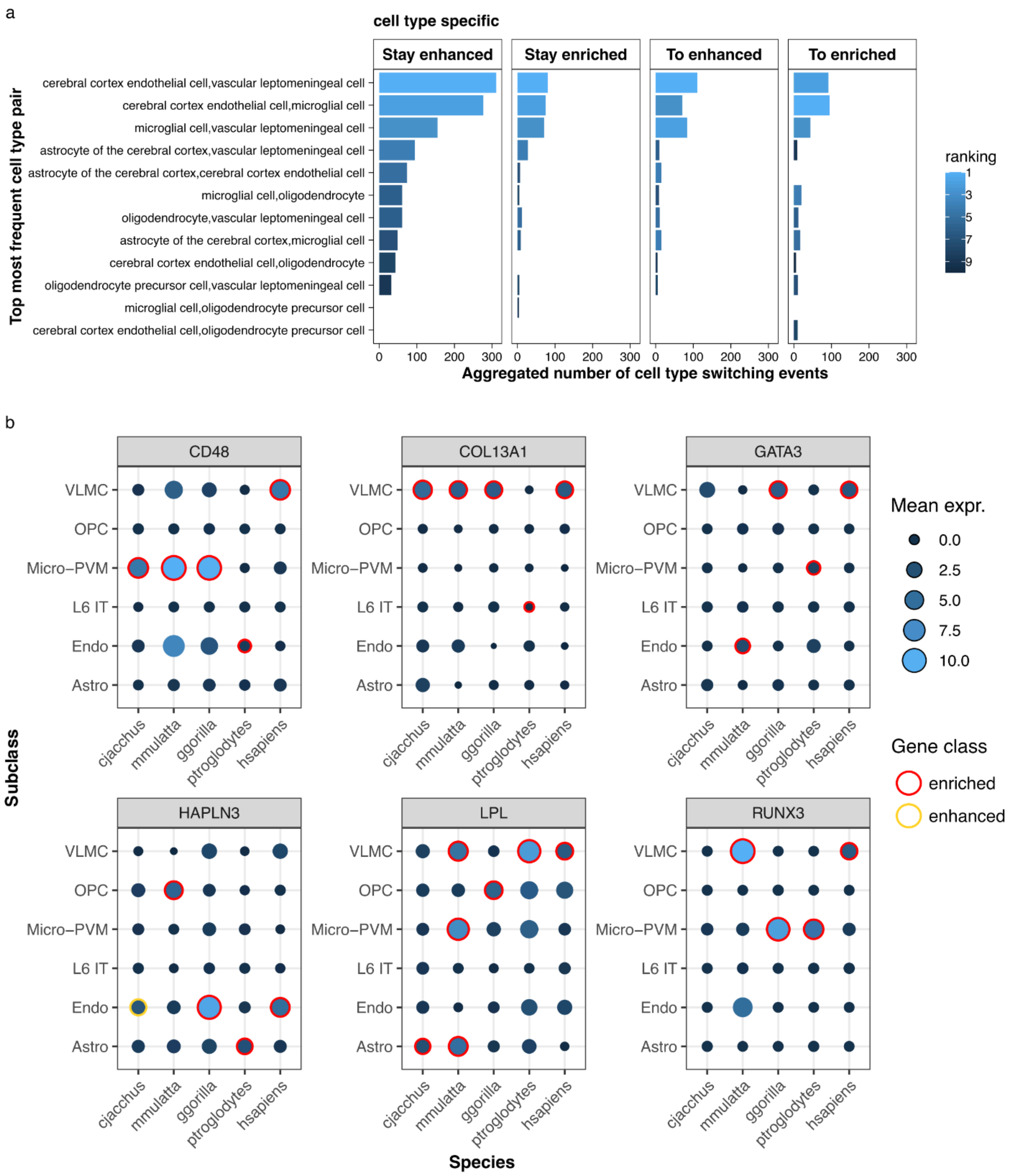

**Supplementary figure 6 Gene specificity class switching mostly happen between transcriptomically similar cell types.** (a) Showing the top cell type pairs which cell type-specific genes switched between them. (b) Showing examples of cell type enriched genes that switched cell type between species with top frequency. Mean expr., mean scaled expression. VLMC, vascular leptomeningeal cell; OPC, oligodendrocyte precursor cell; Micro-PVM, microglial cell; L6 IT, L6 intratelencephalic neurons; Endo, cerebral cortex endothelial cell; Astro, astrocyte of the cerebral cortex.

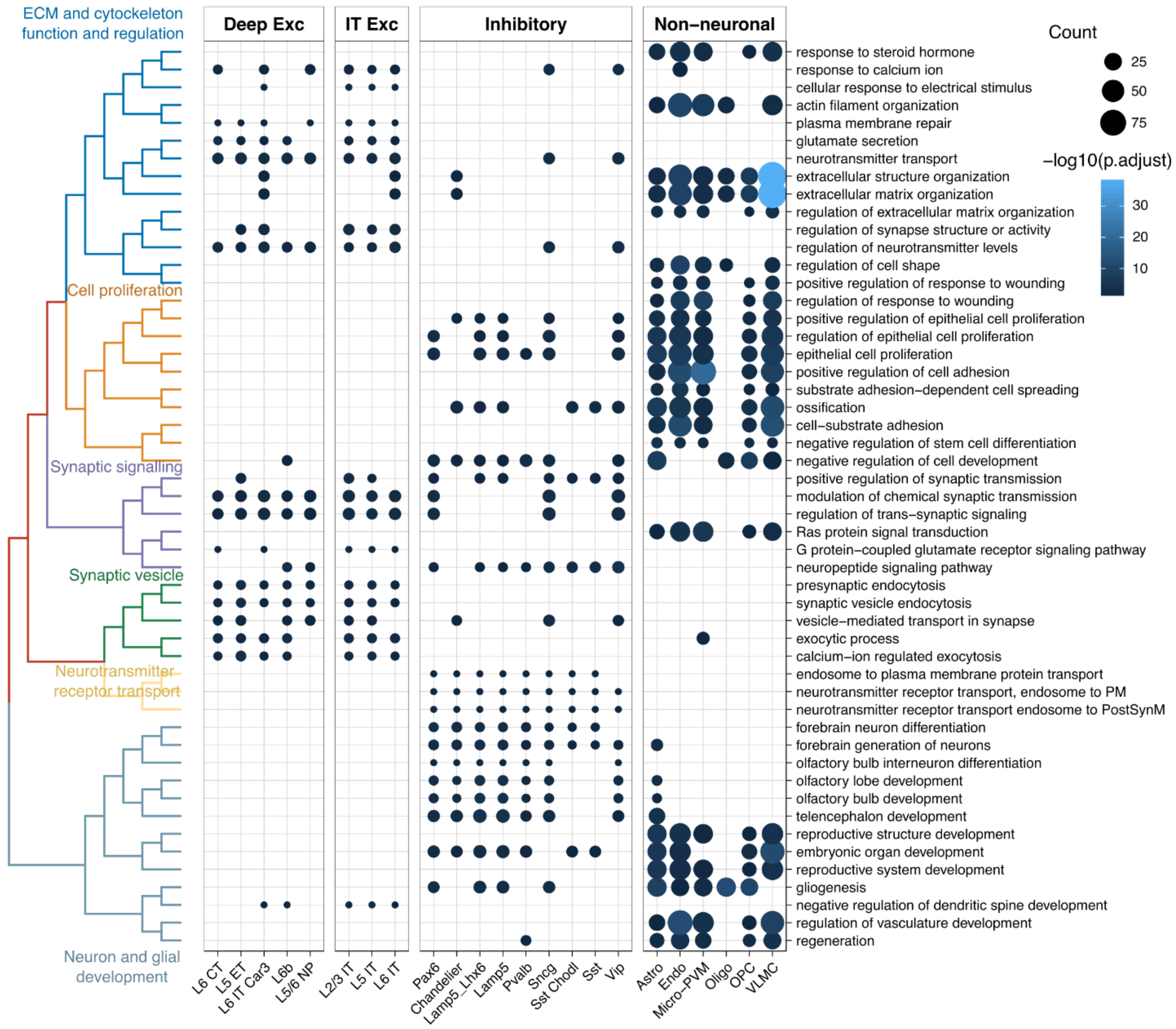


**Supplementary figure 7 GO BP enrichment of genes that stayed in the same specificity class.** Here shows the GO BP enrichment of genes that stayed enriched in the same cell type in any pair of species. Top most frequent enriched terms across each broad type were selected and plotted in the figure. Semantic similarity between the selected GO BP terms were calculated and used for hierarchical clustering, showing on the left and summarised by keywords.

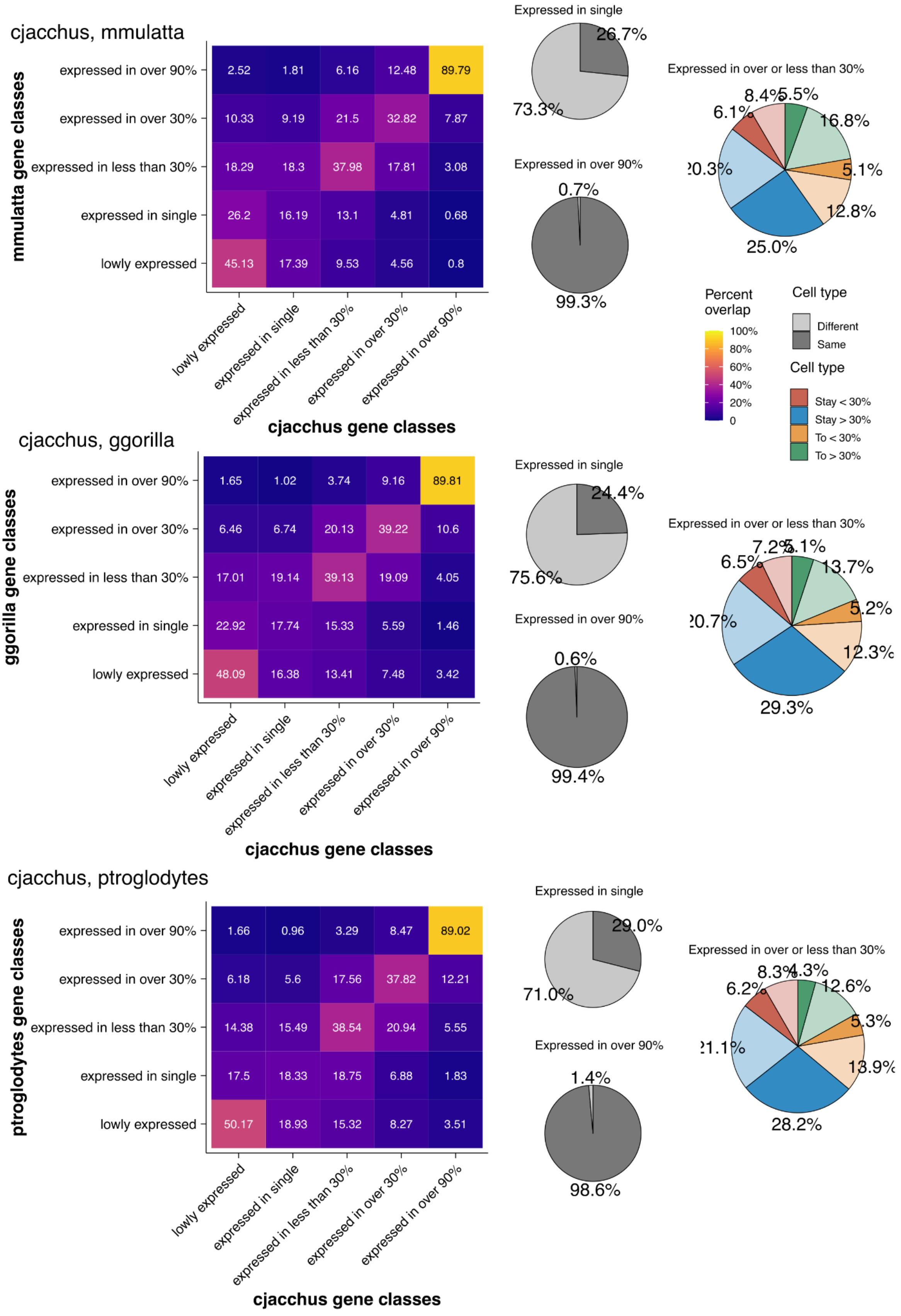

cjacchus, mmulatta
mmulatta gene classes
expressed in over 90%
expressed in over 30%
expressed in less than 30%
expressed in single
lowly expressed
2.52
1.81
6.16
12.48
89.79
10.33
9.19
21.5
32.82
7.87
18.29
18.3
37.98
17.81
3.08
26.2
16.19
13.1
4.81
0.68
45.13
17.39
9.53
4.56
0.8
lowly expressed
expressed in single
expressed in less than 30%
expressed in over 30%
expressed in over 90%
cjacchus gene classes
Expressed in single
26.7%
73.3%
Expressed in over 90%
0.7%
99.3%
Expressed in over or less than 30%
8.4%
5.5%
6.1%
16.8%
5.1%
12.8%
25.0%
Percent overlap
100%
80%
60%
40%
20%
0
Cell type
Different
Same
Cell type
Stay < 30%
Stay > 30%
To < 30%
To > 30%
cjacchus, ggorilla
ggorilla gene classes
expressed in over 90%
expressed in over 30%
expressed in less than 30%
expressed in single
lowly expressed
1.65
1.02
3.74
9.16
89.81
6.46
6.74
20.13
39.22
10.6
17.01
19.14
39.13
19.09
4.05
22.92
17.74
15.33
5.59
1.46
48.09
16.38
13.41
7.48
3.42
lowly expressed
expressed in single
expressed in less than 30%
expressed in over 30%
expressed in over 90%
cjacchus gene classes
Expressed in single
24.4%
75.6%
Expressed in over 90%
0.6%
99.4%
Expressed in over or less than 30%
7.2%
5.1%
6.5%
13.7%
5.2%
12.3%
29.3%
cjacchus, ptroglodytes
ptroglodytes gene classes
expressed in over 90%
expressed in over 30%
expressed in less than 30%
expressed in single
lowly expressed
1.66
0.96
3.29
8.47
89.02
6.18
5.6
17.56
37.82
12.21
14.38
15.49
38.54
20.94
5.55
17.5
18.33
18.75
6.88
1.83
50.17
18.93
15.32
8.27
3.51
lowly expressed
expressed in single
expressed in less than 30%
expressed in over 30%
expressed in over 90%
cjacchus gene classes
Expressed in single
29.0%
71.0%
Expressed in over 90%
1.4%
98.6%
Expressed in over or less than 30%
8.3%
4.3%
6.2%
12.6%
5.3%
13.9%
28.2%

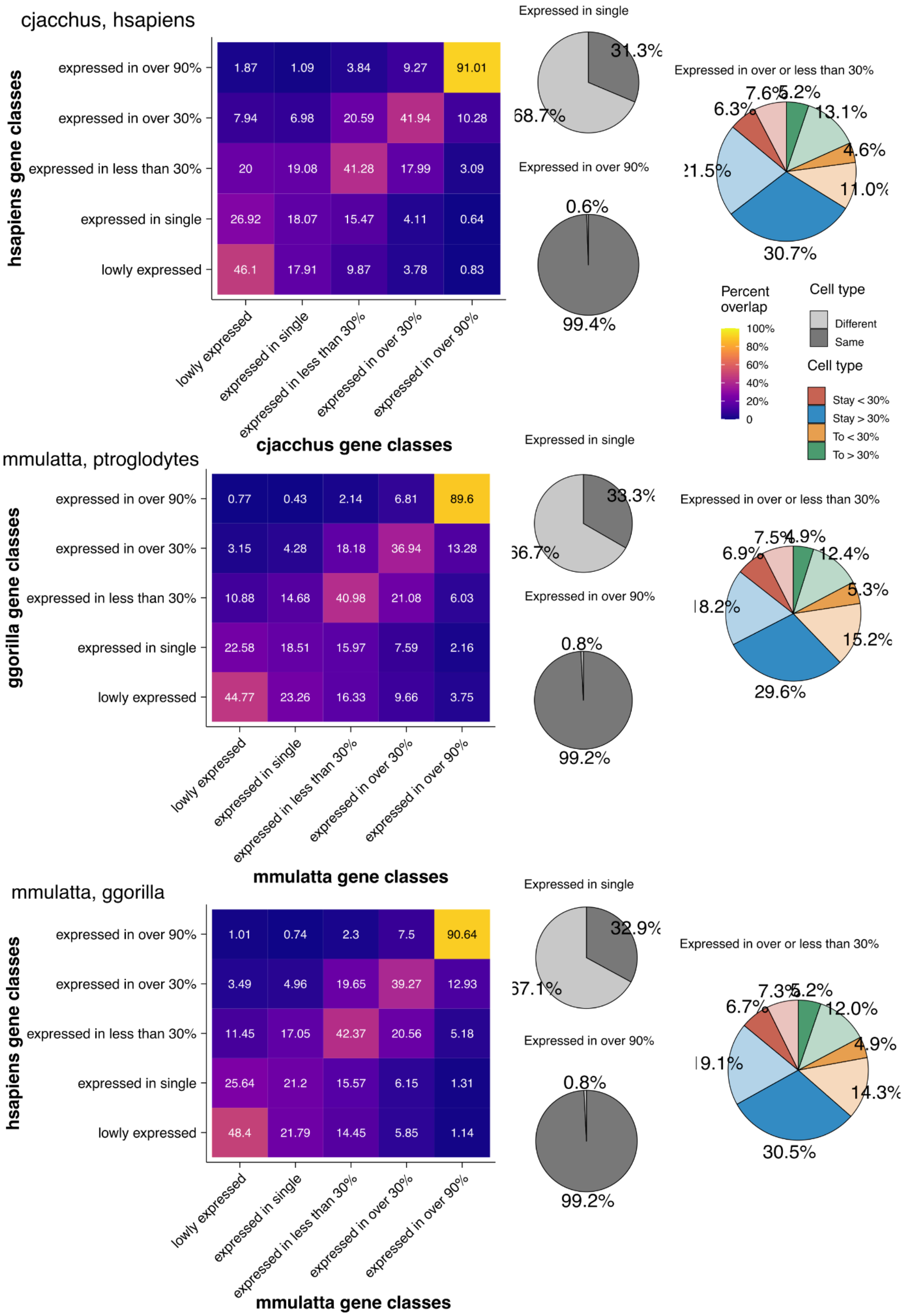
cjacchus, hsapiens
hsapiens gene classes
expressed in over 90%
expressed in over 30%
expressed in less than 30%
expressed in single
lowly expressed
1.87 1.09 3.84 9.27 91.01
7.94 6.98 20.59 41.94 10.28
20 19.08 41.28 17.99 3.09
26.92 18.07 15.47 4.11 0.64
46.1 17.91 9.87 3.78 0.83
lowly expressed
expressed in single
expressed in less than 30%
expressed in over 30%
expressed in over 90%
cjacchus gene classes
Expressed in single
31.3%
68.7%
Expressed in over 90%
0.6%
99.4%
Expressed in over or less than 30%
7.6%
5.2%
6.3%
13.1%
4.6%
21.5%
11.0%
30.7%
Percent overlap
100%
80%
60%
40%
20%
0
Cell type
Different
Same
Cell type
Stay < 30%
Stay > 30%
To < 30%
To > 30%
mmulatta, ptroglodytes
ggorilla gene classes
expressed in over 90%
expressed in over 30%
expressed in less than 30%
expressed in single
lowly expressed
0.77 0.43 2.14 6.81 89.6
3.15 4.28 18.18 36.94 13.28
10.88 14.68 40.98 21.08 6.03
22.58 18.51 15.97 7.59 2.16
44.77 23.26 16.33 9.66 3.75
lowly expressed
expressed in single
expressed in less than 30%
expressed in over 30%
expressed in over 90%
mmulatta gene classes
Expressed in single
33.3%
66.7%
Expressed in over 90%
0.8%
99.2%
Expressed in over or less than 30%
7.5%
4.9%
6.9%
12.4%
5.3%
18.2%
15.2%
29.6%
mmulatta, ggorilla
hsapiens gene classes
expressed in over 90%
expressed in over 30%
expressed in less than 30%
expressed in single
lowly expressed
1.01 0.74 2.3 7.5 90.64
3.49 4.96 19.65 39.27 12.93
11.45 17.05 42.37 20.56 5.18
25.64 21.2 15.57 6.15 1.31
48.4 21.79 14.45 5.85 1.14
lowly expressed
expressed in single
expressed in less than 30%
expressed in over 30%
expressed in over 90%
mmulatta gene classes
Expressed in single
32.9%
67.1%
Expressed in over 90%
0.8%
99.2%
Expressed in over or less than 30%
7.3%
5.2%
6.7%
12.0%
4.9%
19.1%
14.3%
30.5%

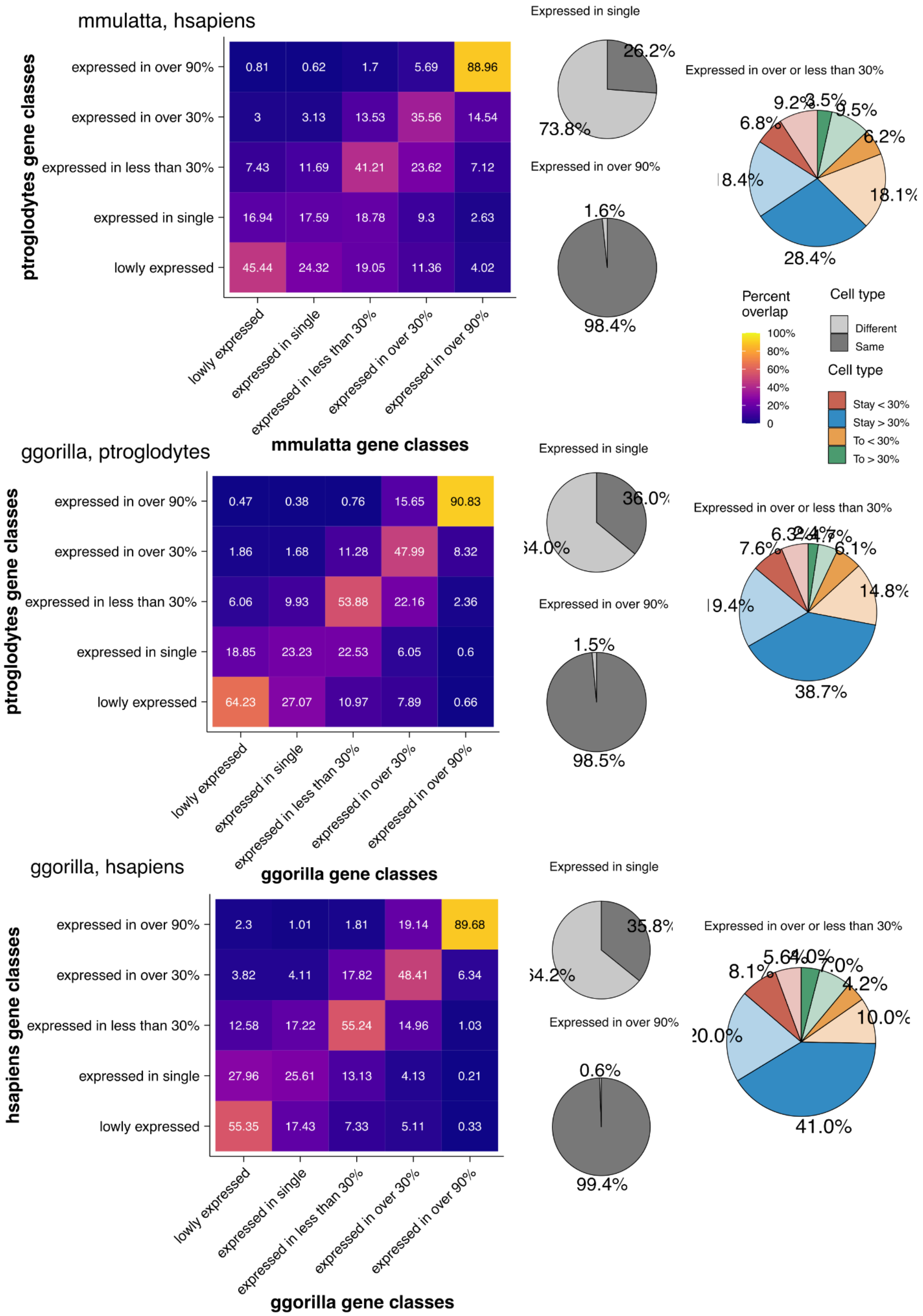

mmulatta, hsapiens
ptroglodytes gene classes
expressed in over 90%
expressed in over 30%
expressed in less than 30%
expressed in single
lowly expressed
0.81 0.62 1.7 5.69 88.96
3 3.13 13.53 35.56 14.54
7.43 11.69 41.21 23.62 7.12
16.94 17.59 18.78 9.3 2.63
45.44 24.32 19.05 11.36 4.02
lowly expressed
expressed in single
expressed in less than 30%
expressed in over 30%
expressed in over 90%
mmulatta gene classes
Expressed in single
26.2%
73.8%
Expressed in over 90%
1.6%
98.4%
Expressed in over or less than 30%
9.2%
3.5%
9.5%
6.8%
6.2%
8.4%
18.1%
28.4%
Percent overlap
100%
80%
60%
40%
20%
0
Cell type
Different
Same
Cell type
Stay < 30%
Stay > 30%
To < 30%
To > 30%
ggorilla, ptroglodytes
ptroglodytes gene classes
expressed in over 90%
expressed in over 30%
expressed in less than 30%
expressed in single
lowly expressed
0.47 0.38 0.76 15.65 90.83
1.86 1.68 11.28 47.99 8.32
6.06 9.93 53.88 22.16 2.36
18.85 23.23 22.53 6.05 0.6
64.23 27.07 10.97 7.89 0.66
lowly expressed
expressed in single
expressed in less than 30%
expressed in over 30%
expressed in over 90%
ggorilla gene classes
Expressed in single
36.0%
Expressed in over 90%
1.5%
98.5%
Expressed in over or less than 30%
7.6%
6.1%
14.8%
38.7%
ggorilla, hsapiens
hsapiens gene classes
expressed in over 90%
expressed in over 30%
expressed in less than 30%
expressed in single
lowly expressed
2.3 1.01 1.81 19.14 89.68
3.82 4.11 17.82 48.41 6.34
12.58 17.22 55.24 14.96 1.03
27.96 25.61 13.13 4.13 0.21
55.35 17.43 7.33 5.11 0.33
lowly expressed
expressed in single
expressed in less than 30%
expressed in over 30%
expressed in over 90%
ggorilla gene classes
Expressed in single
35.8%
Expressed in over 90%
0.6%
99.4%
Expressed in over or less than 30%
8.1%
4.2%
10.0%
41.0%

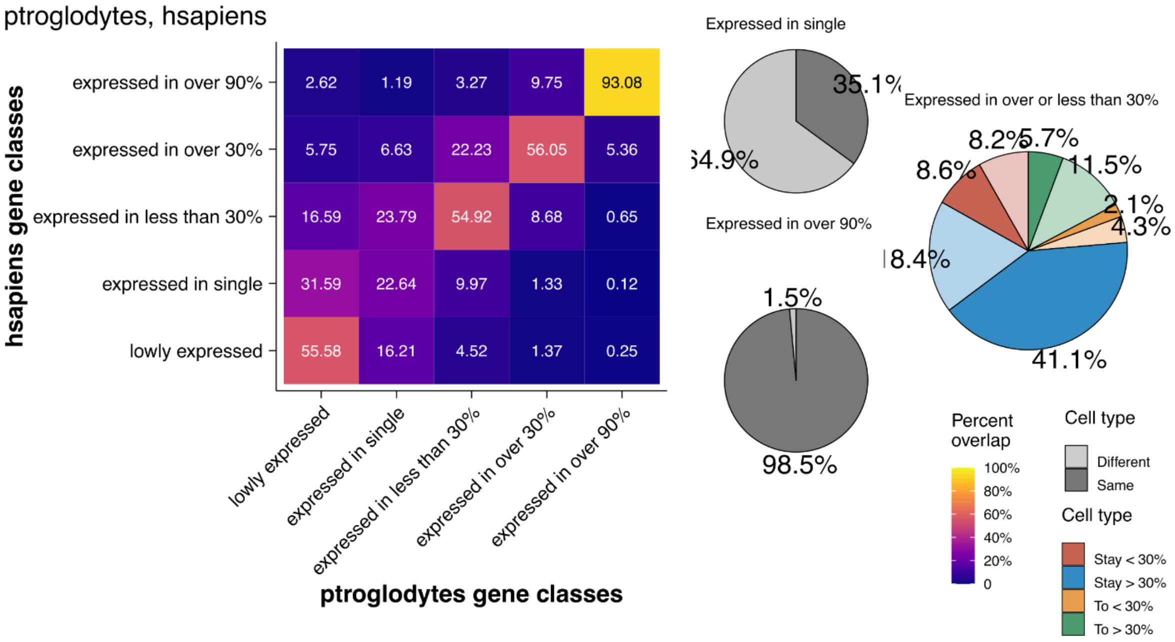


**Supplementary figure 8 Gene distribution class conservation matrix for each species pair.** Showing the average distribution class conservation heatmap (calculation process see methods). Pie chart showing the aggregated percentage of cell type conservation of genes stayed expressed in over 90% of cell types; genes stayed expressed in a single cell type and genes expressed in some cell types.

**a** *H.sapiens* and *P.troglodytes* conserved lowly specific and broadly expressed genes

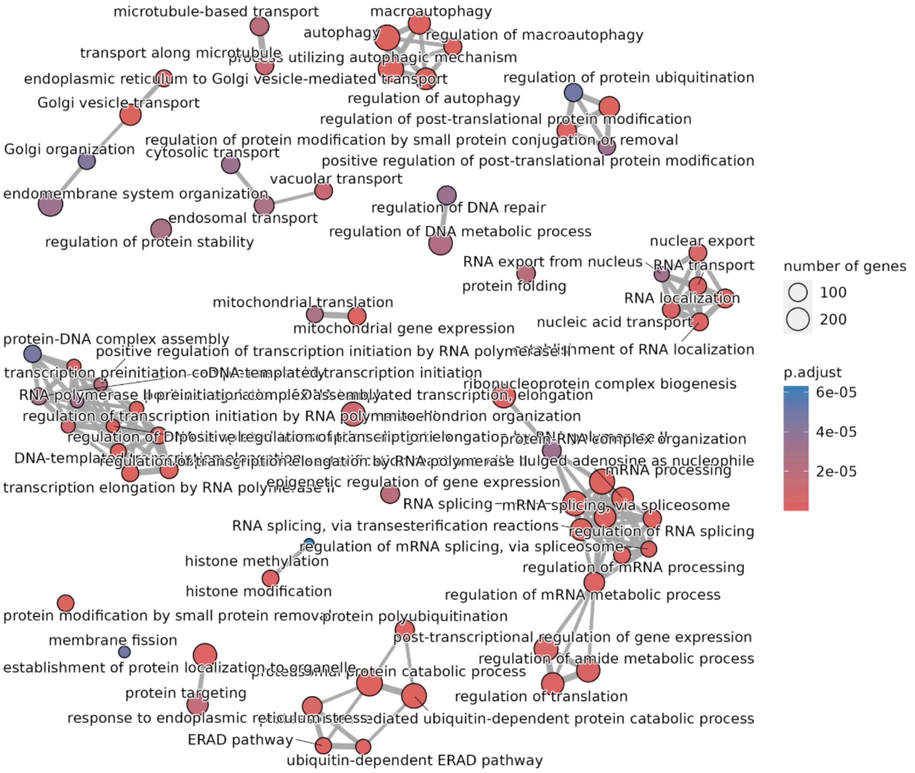


**b** *H.sapiens* and *G.gorilla* conserved lowly specific and broadly expressed genes

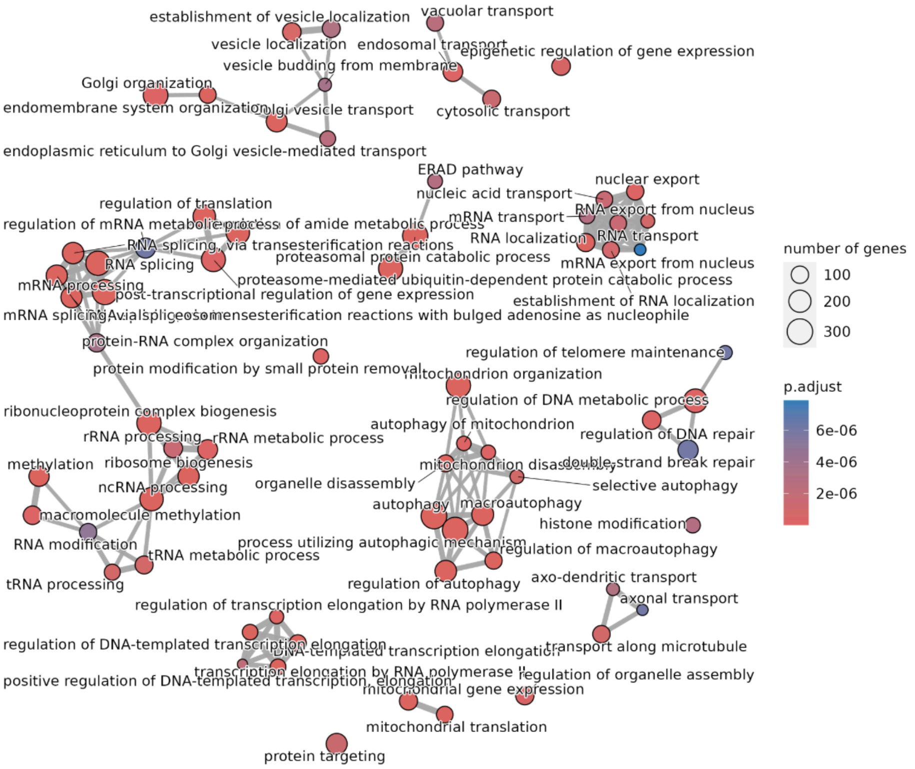

**c** *H.sapiens* and *M.mulatta* conserved lowly specific and broadly expressed genes

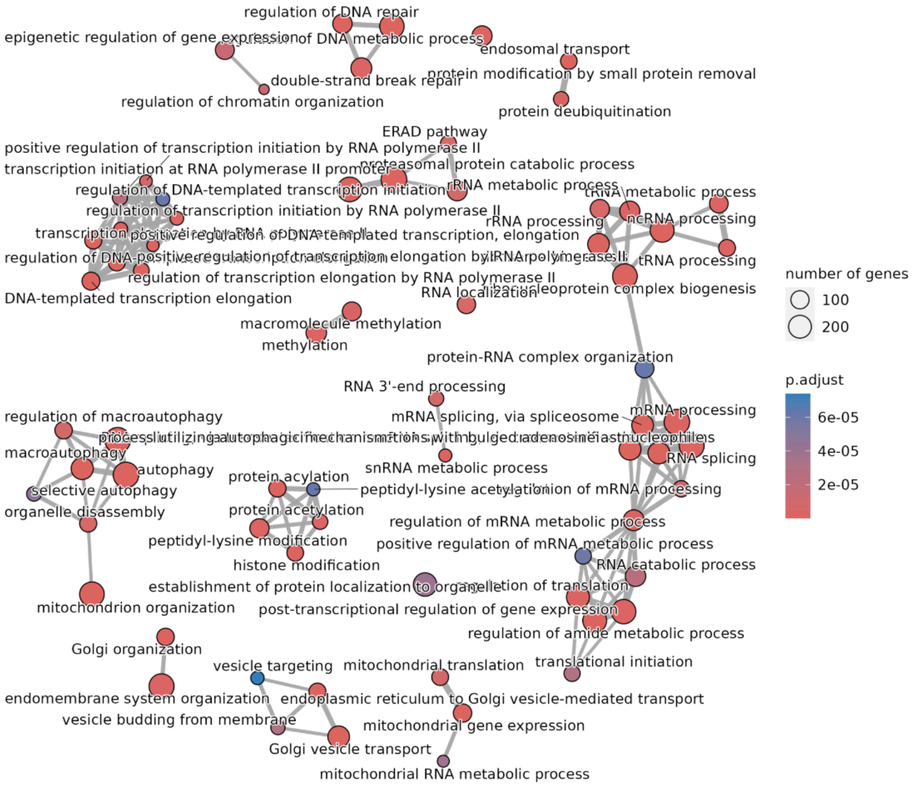


**d** *H.sapiens* and *C.jacchus* conserved lowly specific and broadly expressed genes

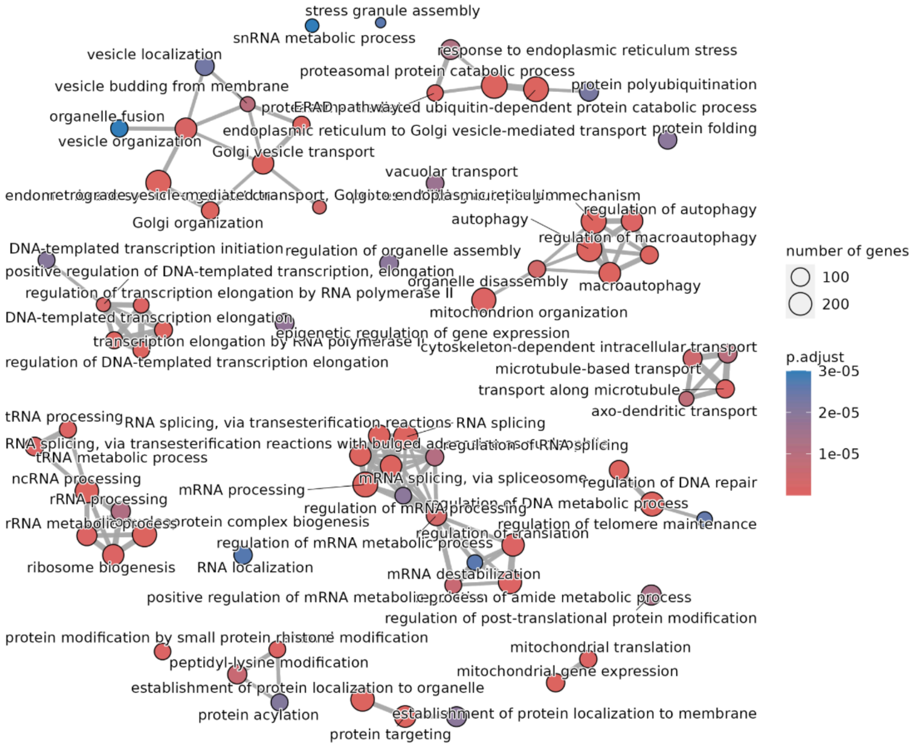


**Supplementary figure 9 Gene ontology enrichment results for genes stayed lowly specific and broadly expressed between human and another species in the primate MTG data.** Figure made with emapplot from the R package enrichplot. Showing 70 categories. P-values

were corrected for multiple comparisons using the Benjamini-Hochberg method, and results were considered significant if P < 0.01.

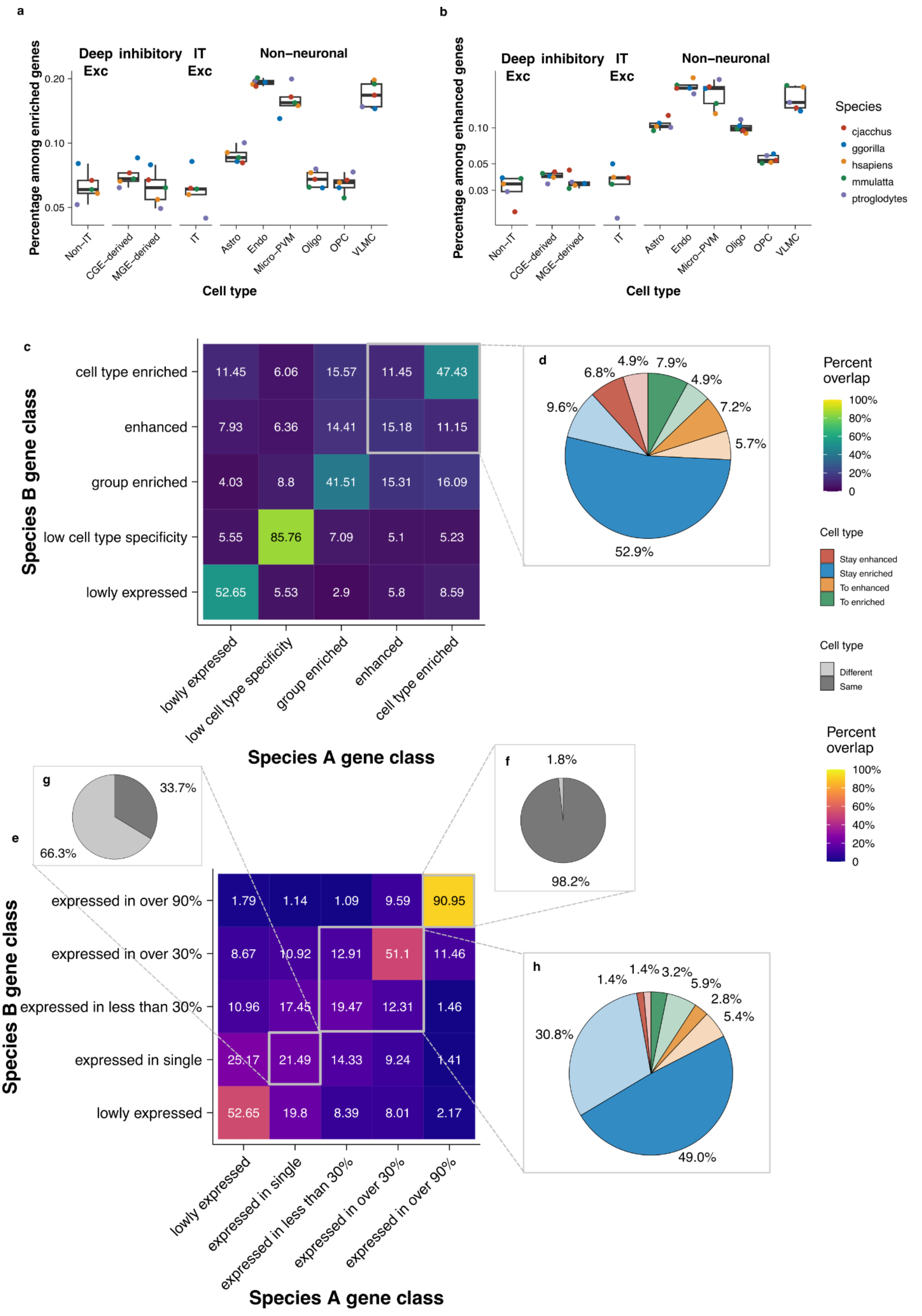

**Supplementary figure 10 Gene class conservation among O2O orthologs in primate MTG dataset using reduced cell type granularity.** Neurons are grouped by reduced granularity but not non-neuronal cells. (a)-(b) Number of cell type-specific genes across species for each cell type. (c) Average specificity class conservation heatmap showing the percent overlap among one-to-one orthologs in various specificity class combinations (calculation process see methods). (d) Pie chart showing the aggregated percentage of cell type conservation and class conservation for genes with cell type level specificity. (e) Average distribution class conservation heatmap (calculation process see methods). (f)-(h) Pie chart showing the aggregated percentage of cell type conservation of genes stayed expressed in (f) over 90\% of cell types; genes stayed expressed in (g) a single cell type and genes expressed in (h) some cell types.

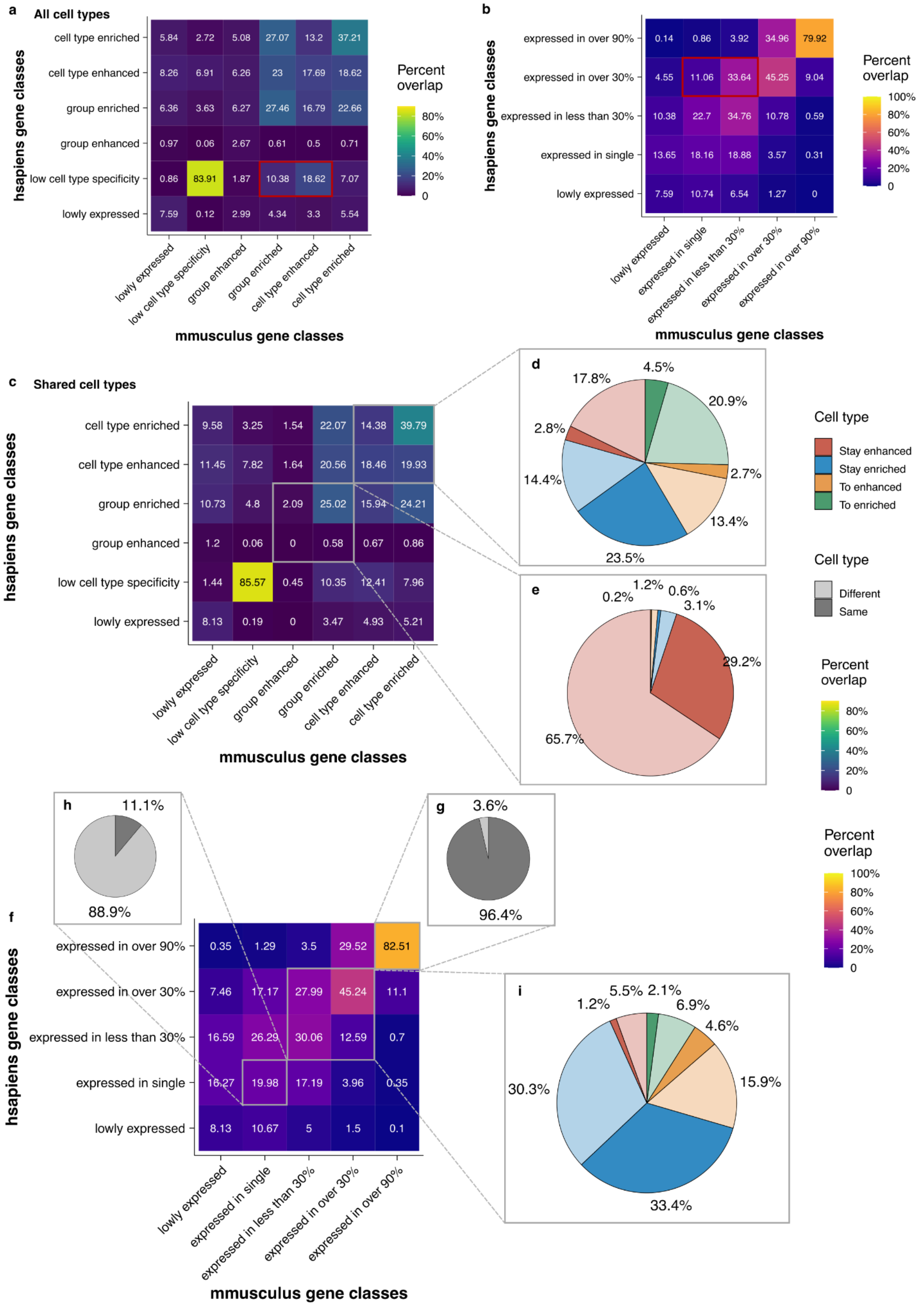


**Supplementary figure 11 Gene class conservation among O2O orthologs in human-mouse bone marrow dataset using all cell types, compared with using only shared cell types.** (a)

Specificity class conservation heatmap showing the percent overlap among one2one orthologs in various specificity class combinations using all cell types (b) distribution class conservation heatmap using all cell types. Red boxes indicate the stripe pattern indicating genes with specificity in mouse but not human, attributed to mouse data-specific cell types. (c) Specificity class conservation heatmap showing the percent overlap among one2one orthologs in various specificity class combinations using shared cell types. (d)-(e) Pie chart showing the aggregated percentage of cell type conservation and class conservation for genes with cell type level specificity (d) or group level specificity (e). (f) Distribution class conservation heatmap using shared cell types. (g)-(i) Pie chart showing the aggregated percentage of cell type conservation of genes stayed expressed in over 90% of cell types (g); genes stayed expressed in a single cell type (h) and genes expressed in some cell types (i).

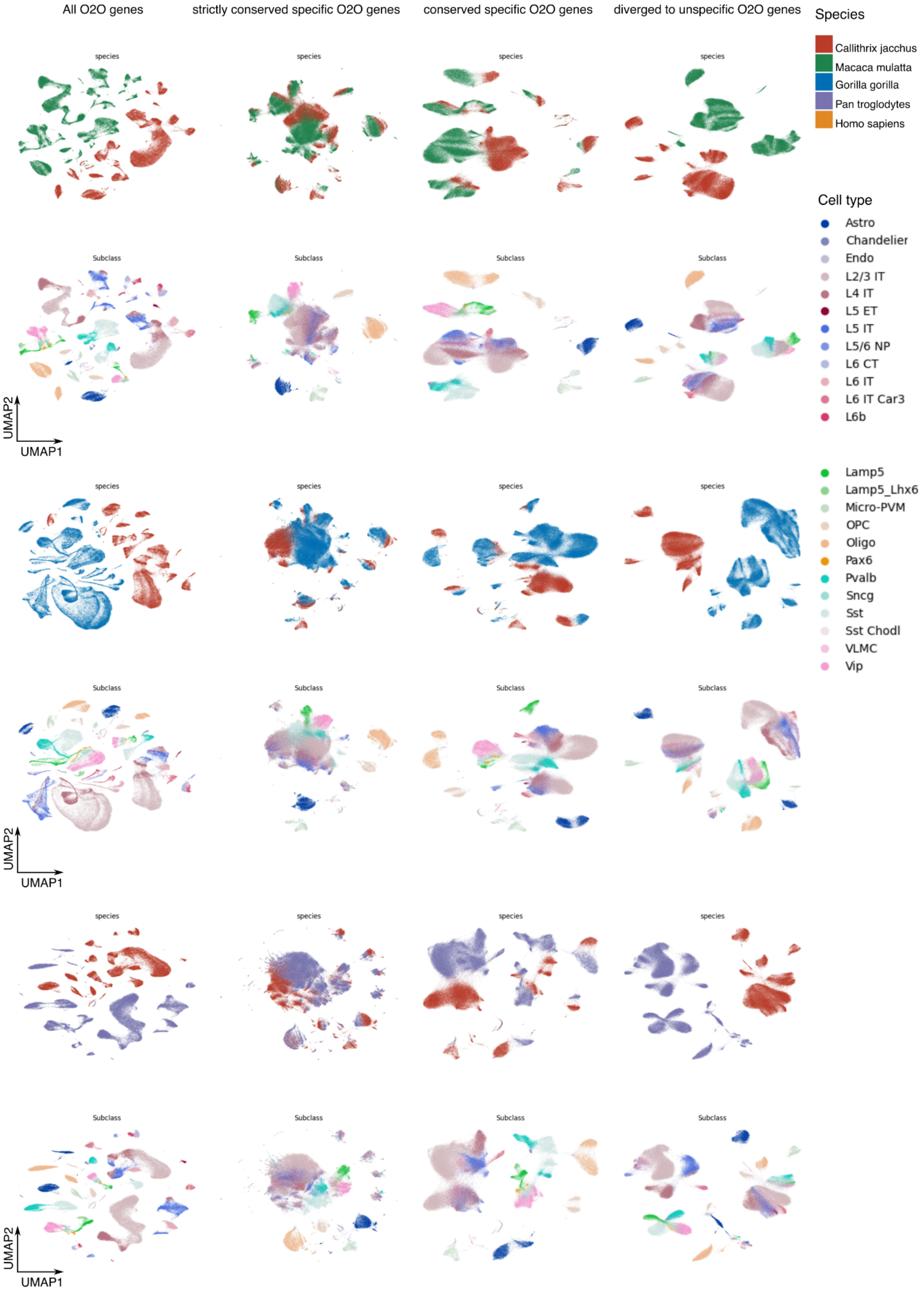

All O2O genes
strictly conserved specific O2O genes
conserved specific O2O genes
diverged to unspecific O2O genes
species
Subclass
UMAP2
UMAP1
Species
Callithrix jacchus
Macaca mulatta
Gorilla gorilla
Pan troglodytes
Homo sapiens
Cell type
Astro
Chandelier
Endo
L2/3 IT
L4 IT
L5 ET
L5 IT
L5/6 NP
L6 CT
L6 IT
L6 IT Car3
L6b
Lamp5
Lamp5_Lhx6
Micro-PVM
OPC
Oligo
Pax6
Pvalb
Sncg
Sst
Sst Chodl
VLMC
Vip

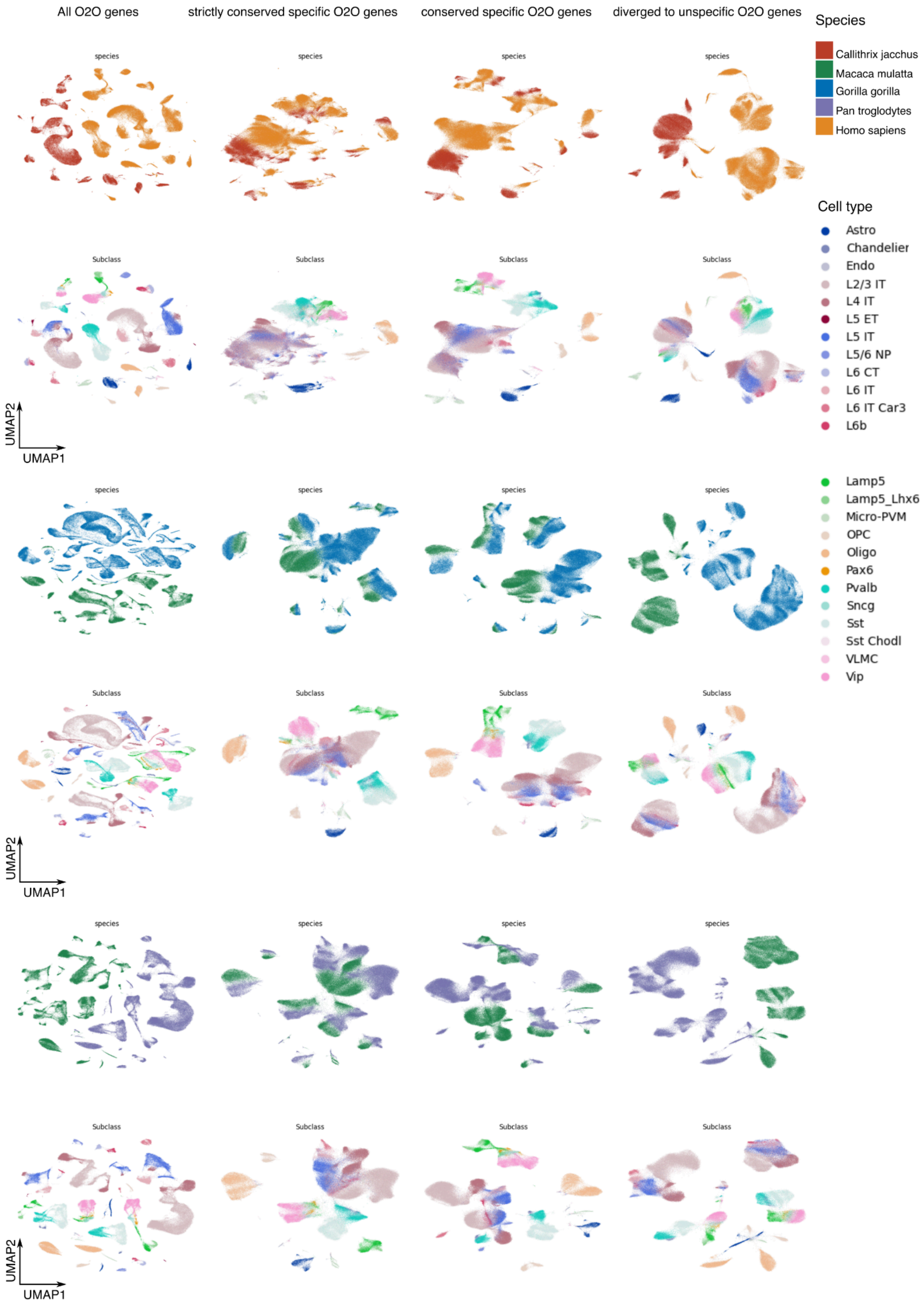
All O2O genes
strictly conserved specific O2O genes
conserved specific O2O genes
diverged to unspecific O2O genes
species
Subclass
UMAP2
UMAP1
Species
Callithrix jacchus
Macaca mulatta
Gorilla gorilla
Pan troglodytes
Homo sapiens
Cell type
Astro
Chandelier
Endo
L2/3 IT
L4 IT
L5 ET
L5 IT
L5/6 NP
L6 CT
L6 IT
L6 IT Car3
L6b
Lamp5
Lamp5_Lhx6
Micro-PVM
OPC
Oligo
Pax6
Pvalb
Sncg
Sst
Sst Chodl
VLMC
Vip

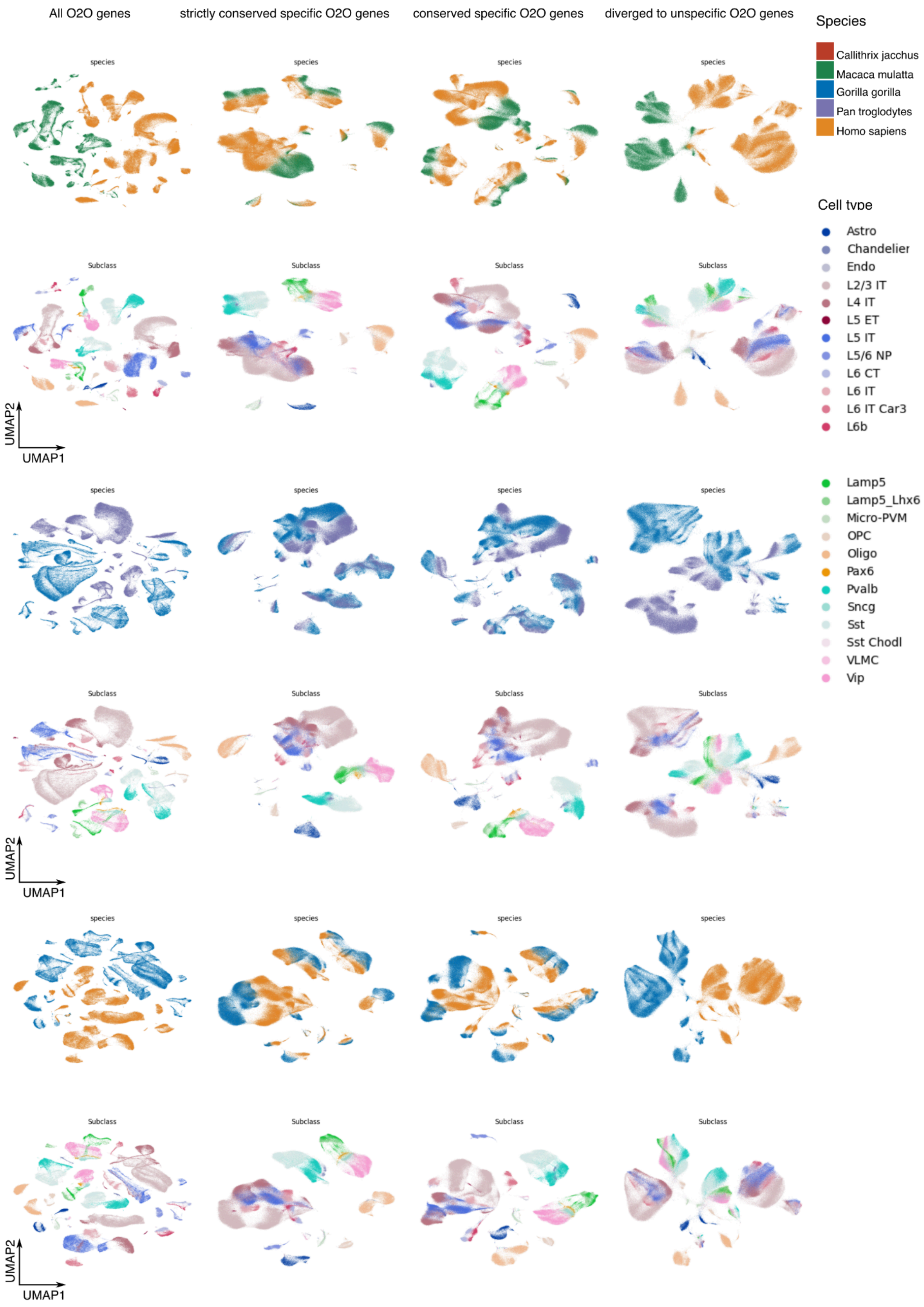


**Supplementary figure 12 Gene class switching explains species effect on transcriptomic space for each species pair.** (a) UMAP visualisation using different sets of O2O orthologs for all species data. Showing the strong species effect using all genes, the reduced species effect using (strictly) conserved specific genes, as well as a purified species effect observed in data using diverged to unspecific genes. O2O, one-to-one; UMAP, Uniform Manifold Approximation and Projection; PC, principal component.

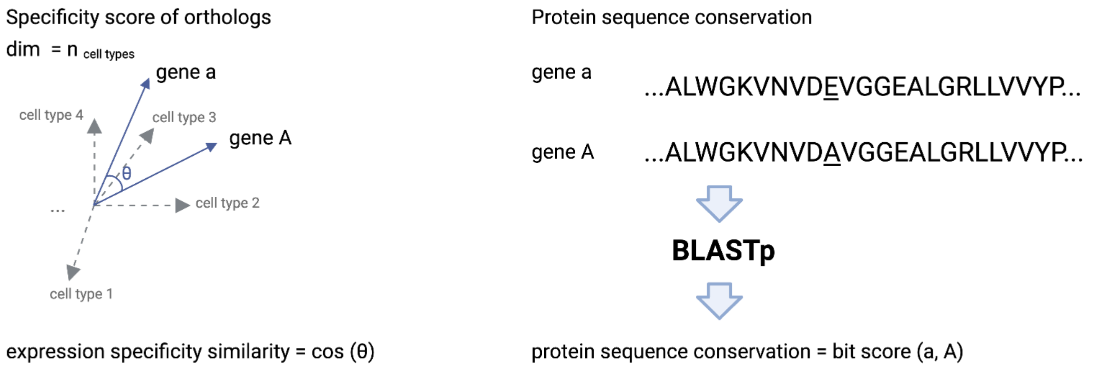


**Supplementary figure 13 Schematic of the comparison of expression specificity similarity and sequence conservation of orthologs.** Gene A and gene a represent orthologs from different species. Expression specificity similarity is calculated with the cosine similarity of gene specificity scores across cell types. Protein sequence conservation is the bit score of BLASTp.

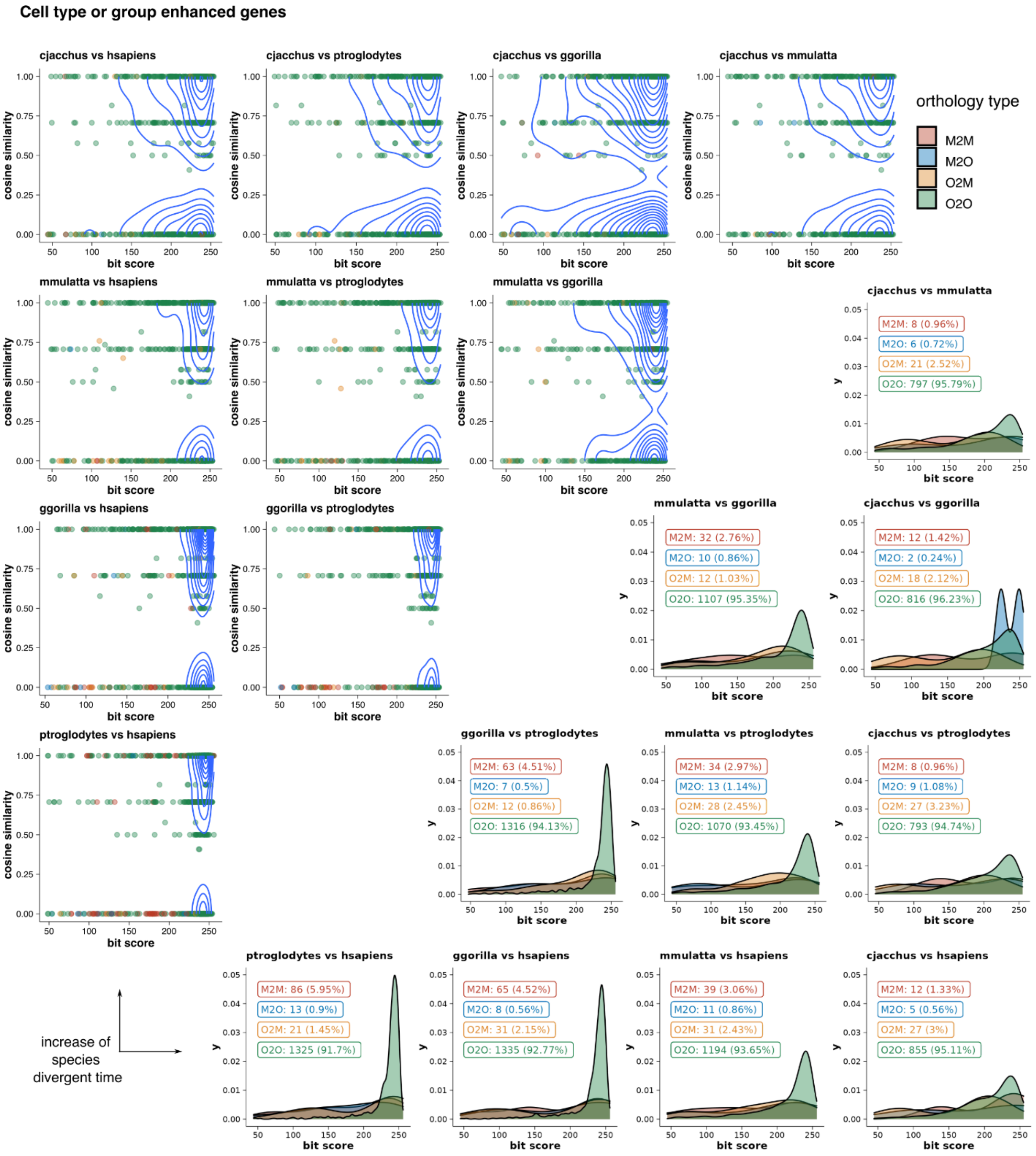


**Supplementary figure 14 Comparing sequence conservation with cell type expression specificity similarity across species distance for enhanced genes.** The top left triangle plots the sequence conservation (bit score) against the expression similarity (cosine similarity) for each pair of cell type enriched orthologous in each pair of species. The bottom right triangle shows the distribution of bit scores for different types of orthologs for the same set of genes. O2O: one-to-one, O2M: one-to-many, M2O: many-to-one, M2M: many-to-many.

**a**

*H.sapiens* and *P.troglodytes* conserved cell type-specific and high sequence conservation genes

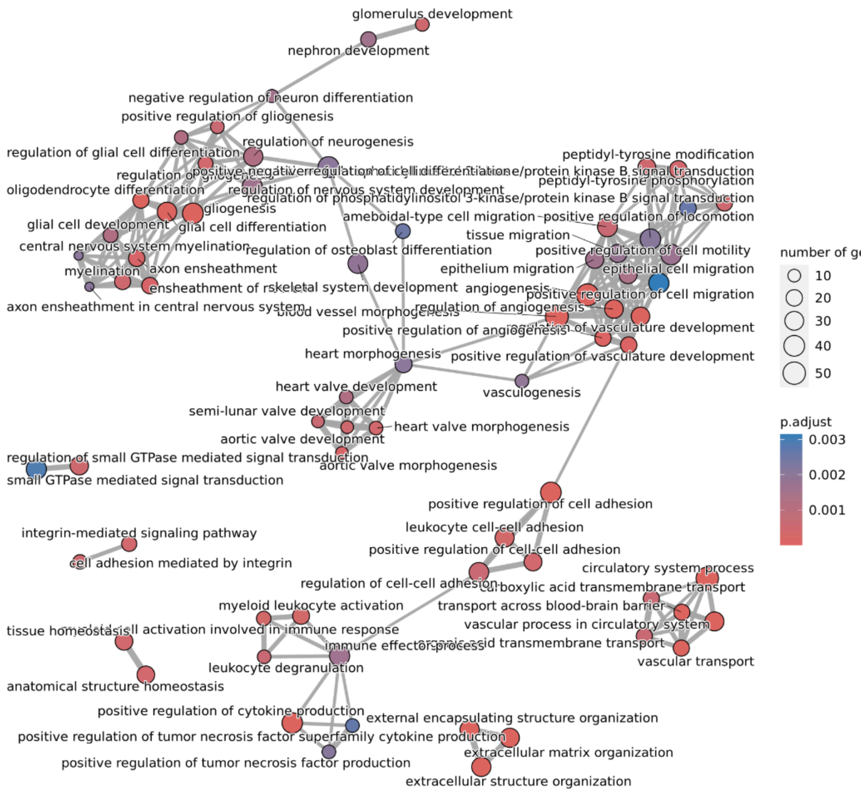


**b**

*H.sapiens* and *G.gorilla* conserved cell type-specific and high sequence conservation genes

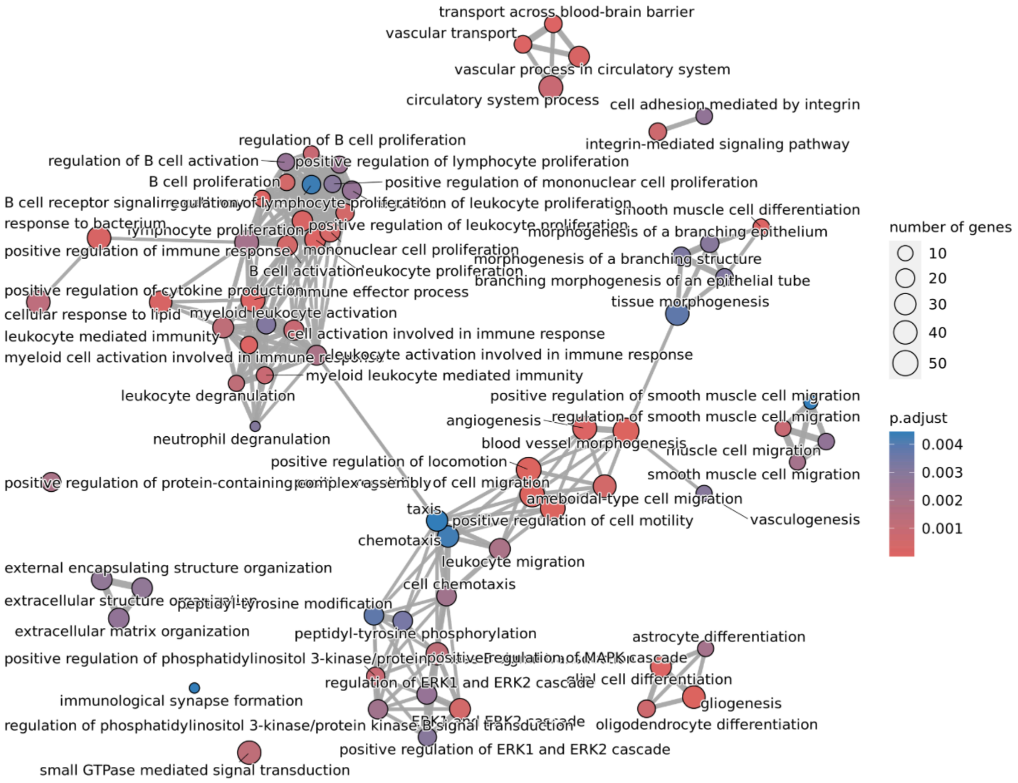

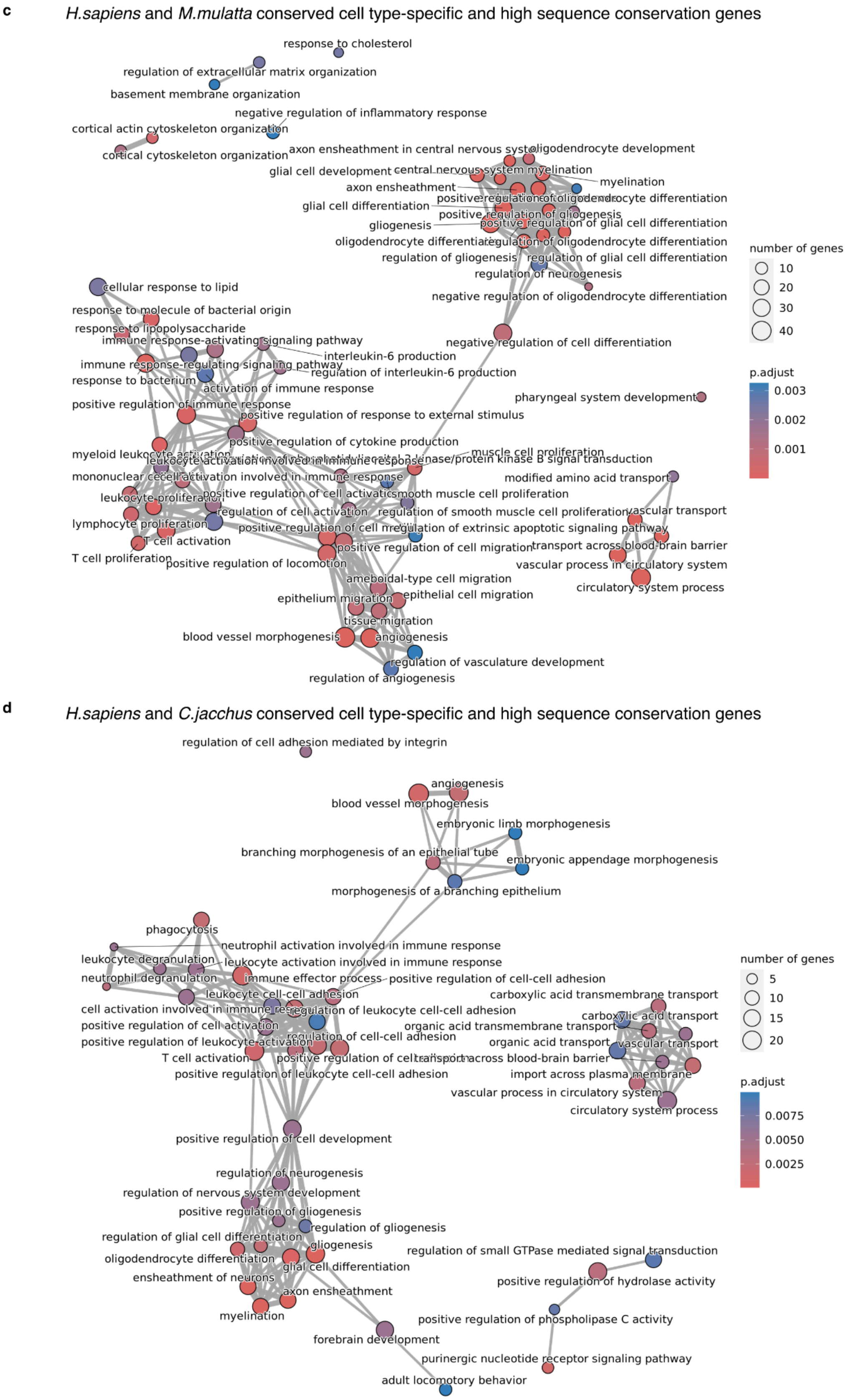


**Supplementary figure 15 Gene ontology enrichment results for genes that have complete specificity conservation and high sequence similarity between human and other species.**

Figure made with emapplot from the R package enrichplot. Showing 70 categories. P-values were corrected for multiple comparisons using the Benjamini-Hochberg method, and results were considered significant if P < 0.01.

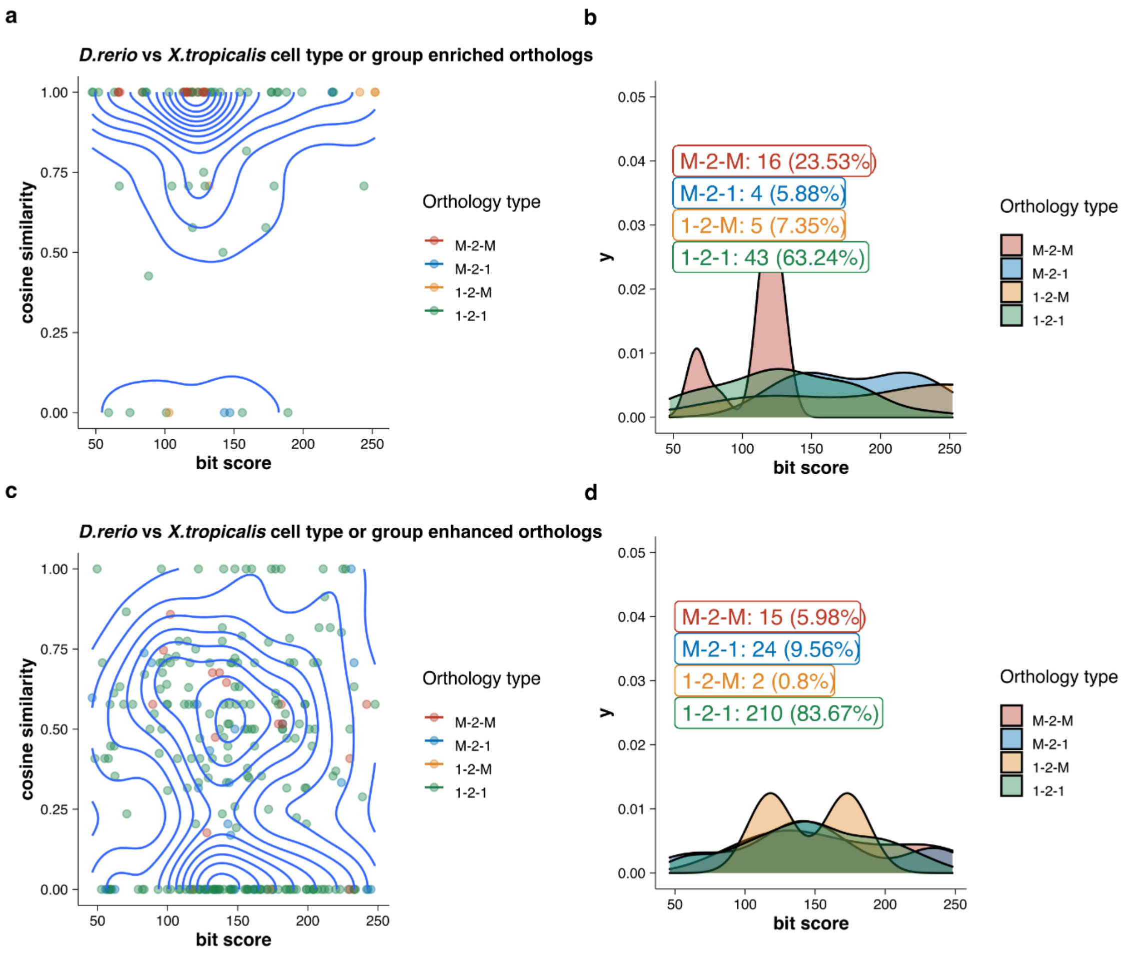


**Supplementary figure 16 Comparing sequence conservation with cell type expression specificity similarity in embryogenesis data.** (a) and (c) plots the sequence conservation (bit score) against the expression similarity (cosine similarity) for each pair of cell type enriched or enhanced orthologous, respectively. (d) and(e) shows the distribution of bit scores for different types of orthologs for the same set of genes. 1-2-1: one-to-one, 1-2-M: one-to-many, M-2-1: many-to-one, M-2-M: many-to-many.
